# Full-length NLRP3 forms oligomeric cages to mediate NLRP3 sensing and activation

**DOI:** 10.1101/2021.09.12.459968

**Authors:** Liudmila Andreeva, Liron David, Shaun Rawson, Chen Shen, Teerithveen Pasricha, Pablo Pelegrin, Hao Wu

**Affiliations:** Department of Biological Chemistry and Molecular Pharmacology, Harvard Medical School, Boston, MA 02115, USA; Program in Cellular and Molecular Medicine, Boston Children’s Hospital, Boston, MA 02115, USA; Harvard Cryo-EM Center for Structural Biology, Boston, MA 02115, USA; Northeastern University, Boston, MA 021115; Instituto Murciano de Investigación Biosanitaria IMIB-Arrixaca, Universidad de Murcia, Murcia, Spain

**Keywords:** NLRP3, NEK7, inflammasome, *trans*-Golgi network, TGN dispersion

## Abstract

The nucleotide-binding domain and leucine-rich-repeat (LRR) containing protein 3 with a pyrin domain (NLRP3) is emerging to be a critical intracellular inflammasome sensor of membrane integrity and a highly important clinical target against chronic inflammation. Here we report that the endogenous, stimulus-responsive form of full-length NLRP3 is a 12-16 mer double ring cage held together by LRR-LRR interactions with the pyrin domains shielded within the assembly to avoid premature activation. Surprisingly, this NLRP3 form is predominantly membrane localized, which is consistent with previously noted localization of NLRP3 at various membrane organelles. Structure-guided mutagenesis reveals that trans-Golgi network dispersion into vesicles, an early event observed for all NLRP3 activating stimuli, requires the double ring cages of NLRP3. Double ring-defective NLRP3 mutants further abolish inflammasome punctum formation, caspase-1 processing and cell death. Thus, unlike other inflammasome sensors that are monomeric when inactive, our data uncover a unique NLRP3 oligomer on membrane that is poised to sense diverse signals to induce inflammasome activation.

---

Inflammasomes are multi-protein complexes that activate inflammatory caspases in response to infection or “damaged-self” signals^1-4^. Activated caspase-1 in turn processes proinflammatory cytokines pro-IL-1β and pro-IL-18 into their mature forms and all inflammatory caspases can cleave GSDMD resulting in pore formation, cytokine release and pyroptotic cell death^3^. The nucleotide-binding domain (NBD) and leucine-rich-repeat (LRR) containing (NLR) protein 3 with a pyrin domain (NLRP3) senses diverse stimuli, from bacterial toxins (e.g. nigericin), extracellular ATP, to particulate particles (e.g. uric acid crystals and amyloids)^5-7^, although none binds to NLRP3 directly. Genetic mutations of NLRP3 cause serious auto-inflammatory disorders, including familial cold autoinflammatory syndrome and Muckle-Wells syndrome^2,8^. Association of NLRP3 with inflammation-driven human diseases, such as cardiovascular diseases, neurological diseases and cancer, further underlies its importance as a clinical target^1,9-11.^

NLR proteins often possess an N-terminal effector domain, a central NACHT ATPase domain that is composed of the NBD, helical domain 1 (HD1), winged helix domain (WHD) and helical domain 2 (HD2), and a C-terminal LRR domain^12^, and for NLRP3, the effector domain is a pyrin domain (PYD) (Fig. 1a). Based on the paradigm established from other NLR proteins such as NLRC4, the inactive state of an NLR is monomeric, and a disk-like oligomer is formed via the NACHT and LRR domains upon inflammasome stimulation^12^. The disk structure then brings together the effector domains to form a template for engaging the apoptosis-associated speck-like protein containing a CARD (ASC) adaptor, which in turn recruits and activates caspase-1. Previously, we used extensive protein engineering to stabilize a monomeric form of PYD-deleted NLRP3 (ΔPYD)^13^. This construct was then used to form a complex with NEK7, the centrosomal serine/threonine kinase that plays a scaffolding function in NLRP3 activation^14-16^, to derive the molecular basis for the NLRP3–NEK7 interaction^13^. However, the physiological form of full-length NLRP3 in its native form remained unknown.

**Fig. 1.**
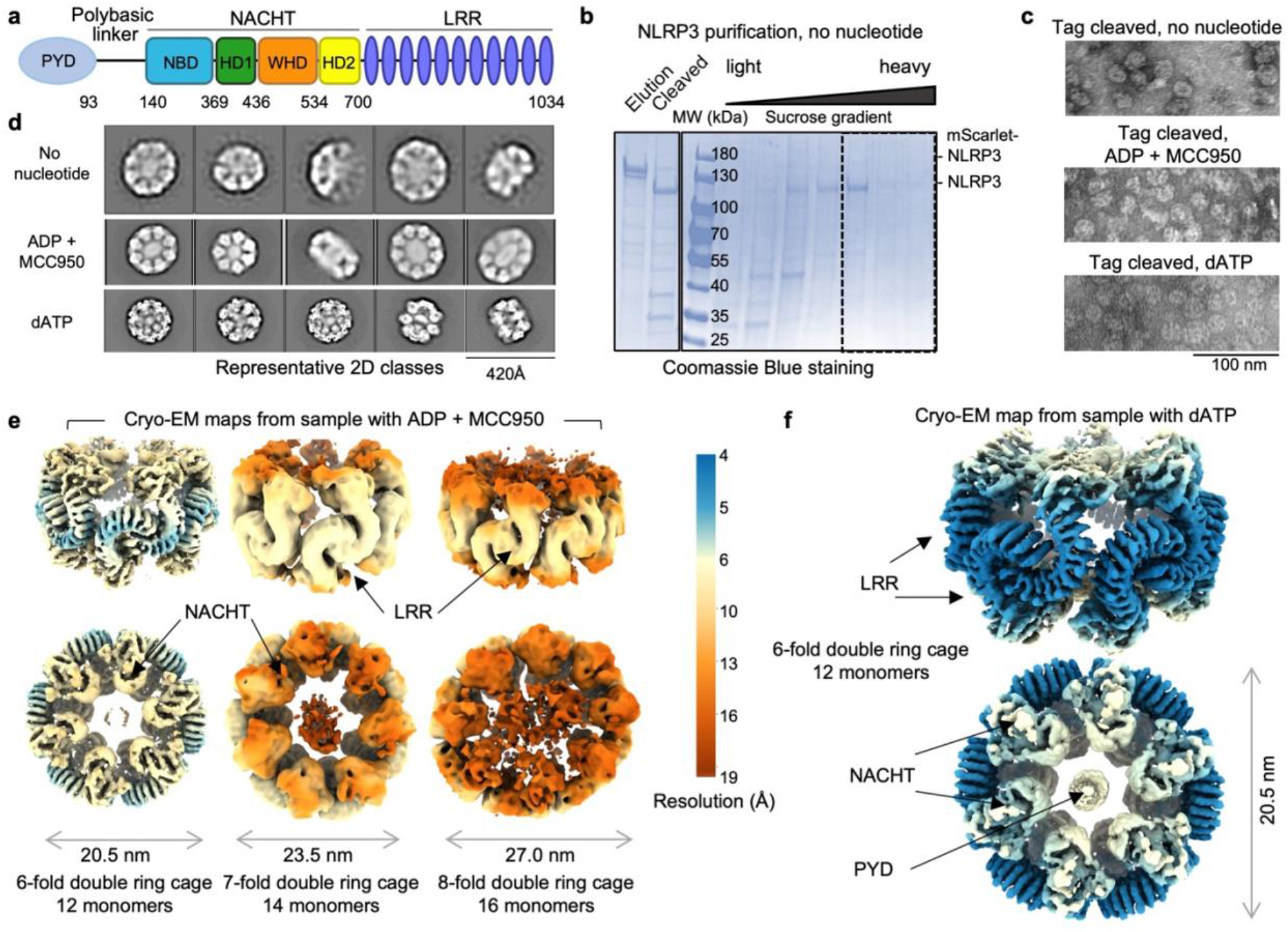
Purification and overall structure of the NLRP3 double ring cage. **a**, Domain organization of NLRP3. **b**, SDS-PAGE of elution and sucrose gradient fractions of NLRP3 purified in the absence of any nucleotide. **c**, Representative negative-staining EM images of NLRP3 oligomers purified without nucleotide (top), with ADP and MCC950 (middle) or with dATP (bottom). **d**, Representative 2D class averages of NLRP3 oligomers purified without any nucleotide (top), with ADP and MCC950 (middle) or with dATP (bottom). **e, f**, Cryo-EM maps of NLRP3 cage species purified in presence of ADP and MCC950 (**e**) or dATP (**f**). Maps are colored by resolution.

In this study, we found that the endogenous form of full-length NLRP3 is not monomeric, but an inactive double ring cage that is primarily membrane localized, which is consistent with previously reported localization of NLRP3 on various membranous organelles – from endoplasmic reticulum (ER), to mitochondria, and to Golgi^17-20^. We showed that this cage structure is required to convert an NLRP3 activating stimulus to trans-Golgi network (TGN) dispersion, an established early event in NLRP3 activation^21,22^. Because dispersed TGN vesicles are further transported to the centrosome (also known as the microtubule-organizing center, or MTOC) to engage the centrosomal NEK7 and form the active NLRP3 inflammasome speck^7,23,24^, our studies uncovered the structural basis for the crucial first step that licenses NLRP3 activation.

## Full-Length NLRP3 exists as large oligomeric structures

To pursue structure determination of full-length NLRP3 in a physiological form, we stably reconstituted it with N-terminal FLAG and mScarlet tags into HEK293T cells which do not express inflammasome proteins. We grew a large amount of these cells in monolayers and lysed them to obtain the cytosolic extract for NLRP3 purification. Anti-FLAG affinity chromatography and FLAG-mScarlet tag removal followed by sucrose gradient revealed large NLRP3 species in heavy sucrose fractions (45-55%) (Fig. 1b). Negative-staining electron microscopy (EM) of the heavy fractions revealed 20-25 nm-sized oligomers of NLRP3 (Fig. 1c), which should be inactive as we did not stimulate the cells expressing them. Since NLRP3 can be stabilized by ADP and the NLRP3 inhibitor MCC950^13^, and dATP was successfully implemented to stabilize the apoptosome^25^ and a plant resistosome^26^, we also performed purifications in the presence of these additives, which resulted in similar NLRP3 particles (Fig. 1c). In addition, the same complexes were observed in the presence of N- or C-terminal tags, indicating that the tags did not interfere with the oligomeric NLRP3 arrangement (Extended Data Fig. 1a). We collected cryo-electron microscopy (cryo-EM) data on these samples. 2D classification of these datasets revealed that dATP gave most homogenous complexes with finer details, as compared to samples in ADP + MCC950 or with no nucleotide (Fig. 1d); an ATPase activity assay showed that dATP, like ATP, was hydrolyzed by NLRP3 (Extended Data Fig. 1b).

## Cryo-EM reveals double ring cage structures of full-length NLRP3

Cryo-EM structure determination by 3D reconstruction uncovered striking NLRP3 double ring cages (Fig. 1e, f, Extended Data Fig. 1c, d, 2 and 3). The ADP and MCC950-containing sample gave rise to 6-, 7- and 8-fold complexes, with 12, 14 and 16 NLRP3 monomers, respectively (Fig. 1e, Extended Data Fig. 1c and 2). These three species are equally distributed indicating heterogeneity in the oligomerization. Despite at relatively low resolutions (5.8 Å, 7.7 Å and 9.5 Å for the 6-, 7- and 8-fold cages, respectively), each NLRP3 monomer with the NACHT and LRR density was clearly recognizable. The 6-fold cage appeared circular from the top view and was refined with the 6-fold symmetry; the 7- and 8-fold cages appeared somewhat elliptical in shape, which disallowed symmetry averaging and thus influenced the resolutions of the complexes (Fig. 1e and Extended Fig. 1c). The NLRP3 sample purified with dATP gave rise to the major class of a 6-fold double ring cage structure at 4.2 Å resolution which was used for model building (Fig. 1f, Extended Data Fig. 1d and 3). Local resolution estimation indicated lower resolutions at the NACHT domains, likely indicative of flexibility, as compared to the better resolved LRRs (Fig. 1e, f, Extended Data Fig. 1c, d). To our surprise, all NLRP3 maps showed presumed density of PYD - the only missing region - inside the NACHT-LRR cage (Fig. 1e, f, Extended Data Fig. 1c, d), though this domain of NLRP3 was least resolved (see below).

## NLRP3 cage formation is facilitated by LRR-LRR and PYD-PYD interactions

The final atomic model contains NACHT and LRR domains, and we did not model the PYD due to the absence of any specific features in its density. The NLRP3 cage is mainly formed by interactions between the curved LRRs in two interfaces, “face-to-face” and “back-to-back” (Fig. 2a-c), with ∼1,200 Å^2^ and ∼500 Å^2^ buried surface areas, respectively, as determined by the PISA server^27^ (Extended Data Fig. 4a). Unlike active apoptosomes and NLRC4 oligomeric rings^28-30^ in which the NACHT domains interact extensively, the NACHT domains in the NLRP3 cage are barely in contact. In addition, they assume an inactive conformation almost identical to those in the autoinhibited NLRC4 and the NLRP3–NEK7 complex structures^13,31^, with only a small, 5º rotation relative to the LRR domain (Extended Data Fig. 4b), which most probably emerges from the “pulling” on the NACHT domains by PYD-PYD interactions.

**Fig. 2.**
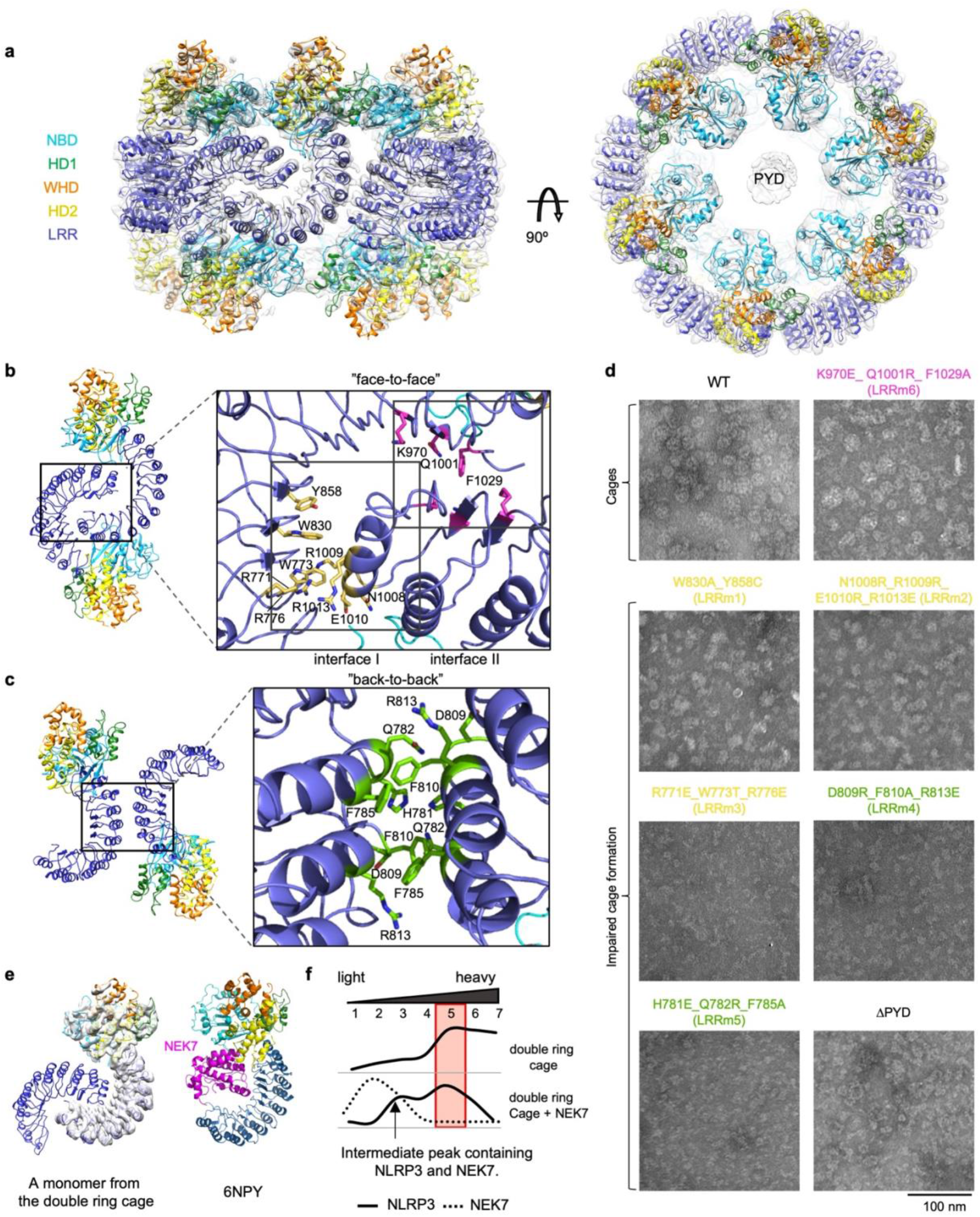
NLRP3 cage is formed by LRR-LRR interactions with PYD inside the cavity. **a**, Atomic model of the NLRP3 cage structure. NLRP3 domains are displayed with the color scheme in Fig. 1a. **b, c**, Overview and detailed views of “face-to-face” (**b**) and “back-to-back” (**c**) interaction interfaces. Residues used for mutagenesis are color-coded: green for “back-to-back” and yellow or magenta for “face-to-face” interfaces. **d**, Representative negative-staining EM images of WT NLRP3 (black) and mutant NLRP3 color coded as in (**b** and **c**). **e**, A comparison of the NLRP3 monomer from the double ring cage with the NLRP3−NEK7 complex (PDB ID: 6NPY)^13^. NEK7 is shown in magenta. **f**, Sucrose gradient profiles of NLRP3 (solid line) and NEK7 (dashed line) in samples containing NLRP3 cage (top) and NLRP3 cage incubated with an excess of NEK7 (bottom) calculated by quantification of SDS-PAGE gels in Extended Data Fig. 5a. The double ring cage containing fractions are highlighted in a red square. An excess of NEK7 disrupts NLRP3 cage.

The larger “face-to-face” interface in the NLRP3 cage is formed by charged and polar residues interacting in agreement with charge complementarity (Fig. 2b, Extended Data Fig. 4c, d). The smaller “back-to-back” interface is less charged, with involvement of many hydrophobic residues (Fig. 2c, Extended Data Fig. 4c, e). Mutational analysis revealed that both interfaces are crucial for the double ring cage formation. Three structure-guided mutants from the “face-to-face” interface (LRRm1: W830A_Y858C; LRRm2: N1008R_R1009R_E1010R_R1013E; and LRRm3: R771E_W773T_R776E) and two from the “back-to-back” interface (LRRm4: D809R_F810A_R813E; and LRRm5: H781E_Q782R_F785A) were impaired in NLRP3 double ring cage formation shown by negative-staining EM (Fig. 2d). Only one mutant, the K970E_ Q1001R_ F1029A mutant (LRRm6) from the “face-to-face” interface, retained the ability to form the double ring cage (Fig. 2d). Surprisingly, deletion of PYD (ΔPYD) was also sufficient to disrupt the NLRP3 double ring structure (Fig. 2d), indicating that PYDs interact and stabilize the complex despite exhibiting only weak density.

The “face-to-face” interface in the double ring cage overlaps with the NEK7-binding surface (Fig. 2e). We thus wondered if NEK7 competes with cage formation. To address this question, we incubated excess recombinant NEK7 with purified NLRP3 cage followed by sucrose gradient. This experiment showed that NEK7 addition resulted in a decrease of the peak for the NLRP3 cage and induced the appearances of smaller NLRP3 species (Fig. 2f, Extended Data Fig. 5a), which suggested that NEK7 competed and partly dissociated the double ring cage. However, NEK7 is a centrosomal kinase and resides predominately on the MTOC shown by immunofluorescence (IF) imaging^23^. Thus, NLRP3 cannot engage NEK7 in resting cells; it colocalized with NEK7 once it was trafficked to the MTOC upon inflammasome stimulation^13,23^, or when the cells were lysed for immunoprecipitation^14-16^. We hypothesize that this spatial separation of NLRP3 and NEK7 in the resting state also offers a mechanism to keep NLRP3 inactive until it is stimulated by pathogenic or damage signals. Of note, NEK7 interaction is not the only requirement for NLRP3 inflammasome formation; a conformational change to an active conformation in the NLRP3 NACHT domain also needs to occur at a certain step after stimulation to allow the assembly of an oligomeric, active NLRP3−NEK7 inflammasome at the MTOC^13^.

## PYDs are shielded within two NACHT-LRR rings to avoid nucleating ASC

Although the NACHT domain conformation of the NLRP3 monomer within the double ring cage suggests that the observed oligomeric complex is inactive (Extended Data Fig. 4b), the extra density (Extended Data Fig. 1c, d) and the role of the PYD in double ring cage formation (Fig. 2d) imply a possibility that the cage contains oligomerized PYD and might be capable of initiating ASC^PYD^ filament assembly. Therefore, we investigated whether NLRP3 cages could induce ASC^PYD^ oligomerization using fluorescence quenching of Alexa488-labeled ASC^PYD^ upon filament formation (Fig. 3a, b, Extended Data Fig. 5b). ASC^PYD^ alone exhibited a low oligomerization rate in the absence of any nucleator. When mixed with NLRP6^PYD-NACHT^ shown previously to nucleate its filament formation^32^, ASC^PYD^ exhibited increased initial rates of fluorescence quenching indicative of increased oligomerization in a manner dependent on the NLRP6^PYD-NACHT^ concentrations (375 nM - 1 µM) (Fig. 3a, b). However, the same concentrations of the NLRP3 cages were unable to nucleate ASC^PYD^ filament formation (Fig. 3a, b), confirming that the double ring cage is inactive not only in its NACHT domain conformation, but also in its ability to nucleate ASC.

**Fig. 3.**
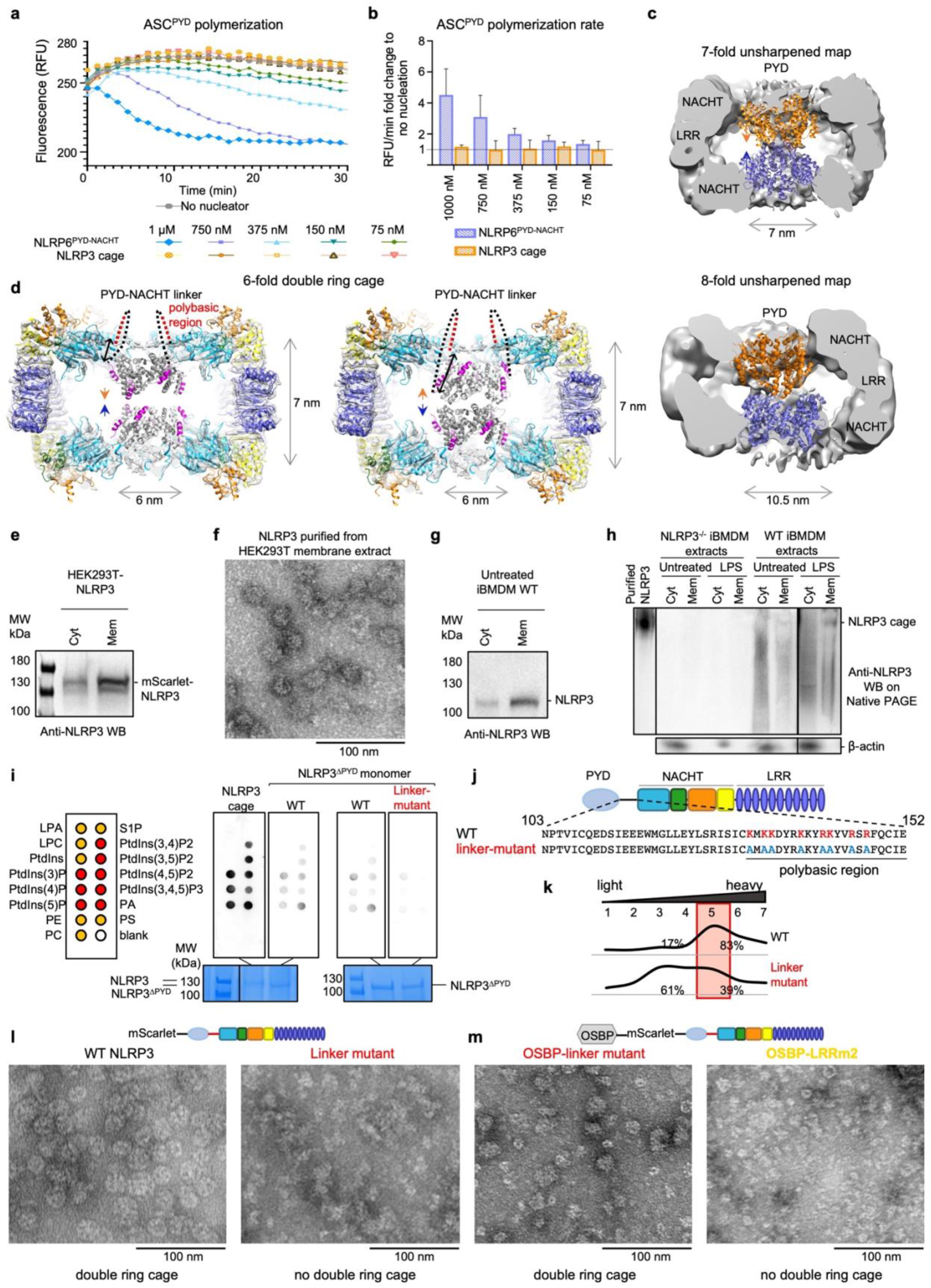
NLRP3 double ring cages are mainly associated with membrane with PYDs shielded within two NACHT-LRR rings to avoid nucleating ASC. **a**, Nucleation of ASC^PYD^ filament formation by NLRP6^PYD-NACHT^ (positive control) or NLRP3 cage, showing fluorescence quenching curves of Alexa488-labeled ASC^PYD^ upon its oligomerization as a function of time (**a**) and the initial rates (**b**). NLRP3 cage did not nucleate ASC^PYD^ filament formation in all concentrations tested. RFU: relative fluorescence unit. Data are presented as mean ± s.d., n=3. **c**, Central slices of the unsharpened maps of 7-(top) and 8-fold (bottom) NLRP3 cages obtained from the ADP + MCC950 sample. The PYD densities inside the NACHT-LRR cages are fitted with short NLRP3^PYD^ helices of 7 or 8 monomers, respectively. The NLRP3^PYD^ filament structure was modelled based on the ASC^PYD^ filament structure (PDB ID: 3J63)^35^. The polymerization directions of the filaments are indicated with the arrows. **d**, A model of a 6-fold NLRP3 cage with two NLRP3^PYD^ helices fitted with the polymerization directions (orange and blue arrowheads) facing the cavity (left). In this orientation, the C-terminus of PYD (magenta) is close to the N-terminus of the NACHT (black double sided arrow). An alternative model with the polymerization directions (orange and blue arrowheads) facing the outside is shown (right). In this orientation, the C-terminus of the PYD is further away from the N-terminus of the NACHT (black double sided arrow). In either model, the PYD to NACHT linker (dotted line) should be exposed at the outside of the cage. Polybasic region of the linker is in red. **e**, Cytosolic (Cyt) and membrane (Mem) extracts of HEK293T cells reconstituted with fluorescently tagged mouse NLRP3 analyzed by WB using anti-NLRP3 antibody. **f**, A negative-staining EM image of NLRP3 oligomers purified from the membrane extract of reconstituted HEK293T cells in the presence of dATP. **g**, Cytosolic (Cyt) and membrane (Mem) extracts of untreated WT iBMDM cells analyzed by WB using anti-NLRP3 antibody. **h**, Native-PAGE of cytosolic and membrane extracts from untreated and LPS-treated WT and NLRP3^-/-^ iBMDMs. The final sample of the NLRP3 cages purified in the presence of dATP for structure determination was used as the control. Anti-NLRP3 and anti-β-actin antibodies were used for the WB. **i**, *In vitro* lipid blot assay of double-ring and monomeric (ΔPYD) forms of NLRP3 with intact (WT) or mutated (“linker-mutant”) polybasic region. On the left, lipid scheme of the membrane with negative-charged (red) and neutral (yellow) lipid head groups is shown. On the bottom, SDS-PAGE is shown as the loading control. **j**, Position and sequence of a WT polybasic region of human NLRP3 and its mutation (linker mutant). Residues involved in TGN binding are colored red and their mutations blue. **k**, Sucrose gradient profiles of WT and linker mutant NLRP3 calculated by quantification of SDS-PAGE gels in Extended Data Fig. 5c. The double ring cage containing fractions are highlighted in a red square. Mutation of the polybasic region impaired double ring cage formation. **l**, Negative-staining EM images of NLRP3 samples from (**k**). Mutation of the polybasic region abolished double ring cage formation. **m**, Negative staining EM images of NLRP3 oligomers purified from OSBP-tagged linker mutant (left) and OSBP-tagged LRRm2 mutant (right). Double ring cage form of NLRP3 was restored by targeting the linker mutant, but not the LRRm2 mutant, back to TGN.

We attempted to fit the presumed PYD density inside the double ring cavity with two short helical models of PYD filaments composed, of 6, 7 or 8 subunits each in the 6-, 7- or 8-fold NLRP3 double ring cage (Fig. 3c, d). There are two potential directions that the short helical segment could fit, one with the C-terminal H6 helices in the PYD oligomer pointing out or pointing in the cavity (Fig. 3d). With the connectivity of the PYD to the NACHT domain, we chose to place the segments with its H6 pointing out to connect more readily to the N-terminus of the NACHT domain. Interestingly, directionality of oligomerization has been observed in the death fold family including for PYD^33-35^, and the fitting we selected would place the elongating ends of the NLRP3 PYD segments for interacting with ASC PYD inside the cavity and facing each other (Fig. 3d). It is tempting to hypothesize that this orientation of the helical PYD segments explains why these cages do not nucleate ASC^PYD^ oligomerization. Together, these observations imply that such a peculiar cage arrangement may serve as a mechanism to shield PYDs within the NACHT-LRR double ring, thus combining oligomerization and inactivity.

## Overexpressed and endogenous NLRP3 are mainly membrane bound

We noticed during the NLRP3 purification that most NLRP3 overexpressed in HEK293T cells was in the membrane pellet rather than the cytosolic extract (Fig. 3e). We thus also performed NLRP3 purification from the membrane pellet after solubilizing it with the n-Dodecyl β-D-maltoside detergent. Visualization of the purified protein using negative-staining EM revealed particles of similar size and shape as observed from the cytosolic extract (Fig. 3f), suggesting that despite not being an integral membrane protein, NLRP3 is mainly membrane bound, perhaps as a peripheral membrane protein. Because the cytosolic purification gave cleaner particles, likely due to less membrane contamination, we did not further pursue structure determination of the NLRP3 particles from the membrane purification.

We also examined the cytosolic versus membrane extracts of endogenous NLRP3 from immortalized bone marrow-derived macrophages (iBMDMs) and found a similar enrichment of NLRP3 in the membrane extract in comparison with the cytosolic extract (Fig. 3g). Since we could not purify endogenous, tag-less NLRP3, we used native PAGE followed by Western blotting to deduce their oligomerization state. We observed that NLRP3 from the iBMDM membrane extract displayed a distinct band that ran at the identical position as purified NLRP3 oligomers, and this band is much more obvious upon LPS priming that increased overall NLRP3 expression (Fig. 3h). By contrast, neither cytosolic nor membrane extract from NLRP3^-/-^ iBMDMs showed signals on the same anti-NLRP3 Western blot. Thus, endogenous NLRP3 from iBMDMs is also mainly membrane bound, and the membrane-localized NLRP3 contains an NLRP3 oligomer identical in size to the purified NLRP3 cage from HEK293T cells. Together, these data support that unlike other inflammasome sensors that are cytosolic and monomeric before activation, NLRP3 forms a physiological cage structure on membranes that may be poised to sense NLRP3-activating signals.

## Membrane association promotes double ring cage formation

To investigate the membrane association of NLRP3, we used a lipid strip assay and found that the purified NLRP3 cage sample bound robustly to mono-, di- and tri-phosphorylated phosphatidylinositols (PtdIns) but not the unphosphorylated PtdIns, as well as phosphatidic acid (PA) that is an acidic precursor of most glycerophospholipids (Fig. 3i). By contrast, monomeric NLRP3 with its PYD removed showed reduced lipid binding, likely due to the loss of avidity in the interaction (Fig. 3i). We further addressed whether membrane association of NLRP3 is crucial for the cage formation using mutagenesis. Since the conserved polybasic region at the PYD to NACHT linker has been shown previously to mediate TGN association (Fig. 3j)^22^, and is predicted to be exposed on the outside of the cage structure (Fig. 3d), we first mutated the conserved polybasic residues to alanine in the monomeric, PYD-deleted NLRP3 (Fig. 3j). In comparison with the WT monomeric NLRP3, the linker mutant displayed an even more reduced binding (Fig. 3i), consistent with the role of these linker residues. We also introduced the same linker mutations to full-length NLRP3. Importantly, although the polybasic region is not at the double ring-forming interfaces, this linker mutant displayed markedly reduced formation of double ring cages, shown by sucrose gradient (Fig. 3k, Extended Data Fig. 5c) and negative-staining EM (Fig. 3l), indicating that membrane binding promotes NLRP3 cage formation.

To address whether the loss of interaction with membranes was responsible for the impaired double ring cage formation in the full-length linker mutant, we targeted it to Golgi as a model membrane source by coupling it to a well described Golgi-binding pleckstrin homology domain of oxysterol-binding protein 1 (OSBP)^36^. Purification of the OSBP-tagged linker mutant (OSBP-linker mutant) revealed the similar double ring cages as for the WT protein under negative-staining EM (Fig. 3m). Although the OSBP-linker mutant sample appeared less homogeneous in size than the WT NLRP3, it contrasted with the linker mutant without OSBP for which mainly random aggregation was apparent (Fig. 3l). We also generated an OSBP-tagged LRRm2 mutant and found that the mutant remained defective in cage formation (Fig. 3m), indicating that the membrane localization per se is not sufficient and that the cage-forming interfaces are also required. We do not know whether a specific, or any membrane localization promotes cage formation. Nonetheless, these data suggest that membranes may serve as a scaffolding platform to promote NLRP3 cage formation. Of note, OSBP tagging restored NLRP3 inflammasome activation of the linker mutant, but not the LRRm2 mutant (see below).

## NLRP3 double ring cage formation is essential for TGN dispersion and NLRP3 activation

Because our NLRP3 cage is the inactive NLRP3 form before activation, we tested its role in TGN dispersion into small vesicles, an early crucial event previously observed for diverse NLRP3 stimuli and viral infection^21,22^. In WT iBMDMs, but not NLRP3^-/-^ iBMDMs, nigericin treatment induced the dispersion of intact TGN from its perinuclear cap-like morphology shown by TGN38 immunofluorescence, and generated NLRP3 inflammasome puncta at the MTOC (Fig. 4a, b, Extended Data Fig. 6a). This effect was validated by another NLRP3 antibody together with an anti-58K Golgi antibody in WT and NLRP3^-/-^ iBMDMs (Extended Data Fig. 6b-d), and channel controls (Extended Data Fig. 6e-g), although residual undispersed 58K Golgi staining remained upon nigericin stimulation due to its localization at both cis- and trans-Golgi. Stable expression of N-terminal mScarlet-tagged NLRP3 in NLRP3^-/-^ iBMDMs rescued TGN dispersion upon nigericin treatment (Fig. 4c). Importantly, treatment with the NLRP3 inhibitor MCC950^37-39^ prior to nigericin stimulation did not affect TGN dispersion while blocking NLRP3 activation (Fig. 4d, Extended Data Fig. 6h-j). These experiments confirmed that TGN dispersion is an upstream event in NLRP3 activation rather than a consequence of NLRP3 activation and pyroptosis. Moreover, the NLRP3 inflammasome puncta formed upon activation co-localized with the TGN38 marker (Fig. 4a, c) – an effect also observed previously that is likely due to trafficking of TGN vesicles to the MTOC^23^.

**Fig. 4.**
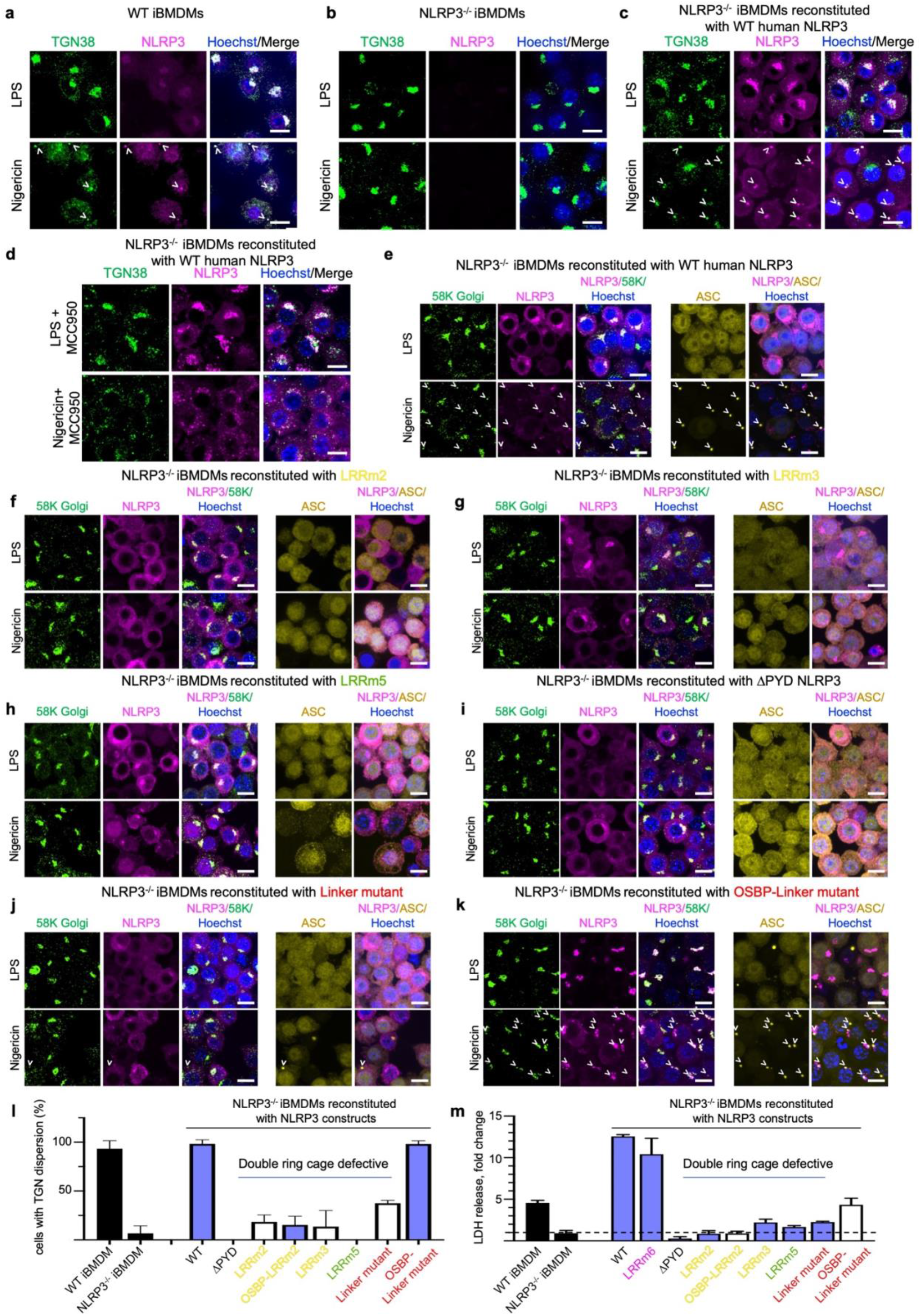
NLRP3 cage is required for TGN dispersion and NLRP3 activation. **a, b**, Confocal imaging of WT (**a**) and NLRP3^-/-^ (**b**) iBMDMs primed with LPS (top) or also treated with 20 µM nigericin for 1 h (bottom) by IF with goat anti-NLRP3 (magenta) and rabbit anti-TGN38 (green) antibodies, and DNA (Hoechst dye, blue). NLRP3 inflammasome specks are labelled with arrowheads. TGN dispersion requires NLRP3. **c**, Confocal imaging of NLRP3^-/-^ iBMDMs reconstituted with WT human mScarlet-NLRP3 primed with LPS (top) or also treated with 20 µM nigericin for 1 h (bottom) for TGN38 (IF, green), NLRP3 (mScarlet, magenta) and DNA (Hoechst dye, blue). These reconstituted cells induced TGN dispersion upon nigericin stimulation identical to WT iBMDMs. **d**, Confocal imaging of NLRP3^-/-^ iBMDMs reconstituted with WT human mScarlet-NLRP3 treated with LPS and MCC950 (top) or pre-treated with MCC950 and treated with 20 μM nigericin for 1 h (bottom) for TGN38 (IF, green), NLRP3 (mScarlet, magenta) and DNA (Hoechst dye, blue). **e-i**, Confocal imaging of NLRP3^-/-^ iBMDMs reconstituted with WT (**e**) or double ring cage disrupting mutants of human mScarlet-NLRP3 (LRRm2, 3, 5 and ΔPYD) (**f-i**) primed with LPS (top) or also treated with 20 µM nigericin for 1 h (bottom) for 58K Golgi protein (IF, green), NLRP3 (mScarlet, magenta) and DNA (Hoechst dye, blue). Disruption of the double ring cage abolished TGN dispersion upon nigericin treatment. **j, k**, Confocal imaging of NLRP3^-/-^ iBMDMs reconstituted with NLRP3 linker-mutant (**j**) or OSBP-linker mutant (**k**) primed with LPS (top) or also treated with 20 µM nigericin for 1 h (bottom) for 58K Golgi protein (IF, green), NLRP3 (mScarlet, magenta) and DNA (Hoechst dye, blue). Targeting the linker mutant to the membrane rescued TGN dispersion. All images are maximum intensity Z projections with scale bars of 10 μm. **l**, Quantification of TGN dispersion based on confocal imaging for WT iBMDMs, NLRP3^-/-^ iBMDMs, and NLRP3^-/-^ iBMDMs reconstituted with WT or mutant human mScarlet-NLRP3, with or without the Golgi-binding OSBP domain. Disruption of the NLRP3 double ring cage abolished TGN dispersion upon nigericin treatment. **m**, Cell death indicated by LDH release for WT iBMDMs, NLRP3^-/-^ iBMDMs, and NLRP3^-/-^ iBMDMs reconstituted with WT or mutant NLRP3, with or without the Golgi-binding OSBP domain. The level of LDH release is shown as a fold change between LPS-primed cells treated or not with nigericin. Data are presented as mean ± s.d., n=3. Double ring cage disrupting mutations abolished NLRP3-mediated cell death.

To address the functional role of NLRP3 cage formation in NLRP3 activation, we reconstituted NLRP3^-/-^ iBMDMs with WT mScarlet-NLRP3 and 4 representative cage-disrupting mutants − 2 at the larger “face-to-face” interface (LRRm2 and LRRm3), 1 at the smaller “back-to-back” interface (LRRm5), and 1 with ΔPYD − and imaged the cells for NLRP3, ASC, 58K Golgi and Hoechst upon LPS priming and upon further nigericin stimulation (Fig. 4e-i). Mouse Anti-58K Golgi was used instead of rabbit anti-TGN38 to allow simultaneous staining for ASC, which also utilized a rabbit antibody. In WT NLRP3-reconstituted NLRP3^-/-^ iBMDMs, TGN dispersed upon nigericin treatment, and the NLRP3 and ASC inflammasome specks also contained 58K Golgi (Fig. 4e), further supporting the trafficking of TGN vesicles. In NLRP3^-/-^ iBMDMs reconstituted with the 4 cage-defective mutants, impaired TGN dispersion and defective inflammasome punctum formation were observed upon nigericin treatment (Fig. 4f-i).

We used OSBP-linked NLRP3 linker mutant and LRRm2 mutant to further dissect the role of the NLRP3 cage in NLRP3 activation. While the linker mutant, which was defective in cage formation (Fig. 3l), also failed to cause TGN dispersion and inflammasome punctum formation (Fig. 4j), the OSBP-linker mutant, which was able to form the cage structure (Fig. 3m), mediated TGN dispersion and inflammasome punctum formation upon nigericin stimulation (Fig. 4k). By contrast, OSBP did not rescue cage formation of the LRRm2 mutant (Fig. 3m) and did not induce TGN dispersion and inflammasome punctum formation upon nigericin stimulation (Extended Data Fig. 6k). Thus, the double ring cage structure is required for nigericin-induced TGN dispersion and NLRP3 activation.

Quantified percentages of cells with TGN dispersion upon nigericin stimulation from the microscopy images were 93%, 98 and 98% for WT iBMDMs, NLRP3^-/-^ iBMDMs reconstituted with WT NLRP3, and NLRP3^-/-^ iBMDMs reconstituted with OSBP-linker mutant, respectively (Fig. 4l). 38% of the linker mutant-reconstituted iBMDMs exhibited TGN dispersion, and iBMDMs reconstituted with the remaining mutants showed even less TGN dispersion upon nigericin stimulation (≤ 18%) (Fig. 4l). Further assessment of inflammasome activation from caspase-1 processing by Western blotting (Extended Data Fig. 7a) and cell death by release of the lactate dehydrogenase (LDH) (Fig. 4m) was consistent with the degrees of TGN dispersion. Importantly, at least LRRm2 and LRRm5 retained similar NEK7 interaction as the WT (Extended Data Fig. 7b), further supporting that their defective functions are due to impaired ability to disperse TGN. Thus, TGN dispersion and inflammasome activation upon nigericin treatment is dependent on cage-forming NLRP3.

## Conclusion

In summary, our discovery of a novel double ring cage structure of full-length NLRP3 sheds light on how the NLRP3 activation pathway may be organized. First, at its constitutive state before or after priming but without stimulation, NLRP3 is already mainly localized on membranes, and the membrane association acts as a scaffolding platform to promote the assembly of an inactive cage structure. This observation is consistent with a resting-state oligomer implicated previously by bioluminescence resonance energy transfer (BRET)^40^. Importantly, the observed double ring cage shields the signal transducing PYDs within the assembly and thus prevents premature ASC recruitment. This strategy of using a “cage” to restrict protein activity is a recurring theme in biology, e.g., in the DegP and the proteasome system in which the proteolytic activity is caged inside the oligomers^41,42^. We do not know exactly which membrane these NLRP3 cages reside; however TGN membrane localization of NLRP3 is visible before nigericin stimulation (Fig. 4), and additional NLRP3 may be recruited to the TGN upon nigericin treatment as suggested previously^9^. Because almost all NLRP3 activating stimuli converges to K^+^ efflux^43,44^, it is reasonable that a drop in intracellular K^+^ concentration could enhance charged interactions and TGN recruitment of NLRP3.

Second, all double ring disrupting mutants of NLRP3 were defective in mediating TGN dispersion to stop the pathway at this early event common to all NLRP3 stimulations^9^ (Extended Data Fig. 8a). Though the exact mechanism of the double ring mediated TGN dispersion remains to be discovered, NLRP3 does not require a conformational change in its NACHT domain to support TGN dispersion. Indeed, MCC950, which was shown to block the inflammasome activation by keeping NLRP3 in a “closed” inactive conformation^13,37-39^, did not interfere with nigericin-induced TGN dispersion. During the preparation of this manuscript, a complementary preprint on NLRP3 decamer in complex with MCC950 was released^45^ showing a clear MCC950 density within an NLRP3 cage. Together, these findings suggest that the inactive conformation of NLRP3 within the double ring cage is compatible with its hypothesized role in TGN dispersion (Extended Data Fig. 8b).

Third, we hypothesize that this peculiar NLRP3 arrangement serves as a sensor coupling NLRP3-activating stimuli with NLRP3 transport on TGN vesicles to its activation center at the MTOC^23,24^. Without TGN dispersion, there would have been no NLRP3-containing vesicles to transport, explaining its importance. Thus, by controlling TGN dispersion, the NLRP3 double ring licenses NLRP3 transport and subsequent activation (Extended Data Fig. 8a). In case NLRP3 gets transported to MTOC in the form of the double ring cage, it is tempting to propose that once reaching the MTOC, the resident protein NEK7 might break the NLRP3 double ring on TGN vesicles to enable further conformational rearrangement. We do not know when and where the NACHT domain changes to its active state that is likely needed for oligomerization of the NLRP3– NEK7 complex^13^. One possibility is that the NACHT domain is flexible, supported by the lower resolution of this region, and that the presumed active NACHT conformation is compatible with the double ring structure (Extended Data Fig. 8b). Thus, NEK7, NLRP3, and most likely ASC may all be required to stabilize the inflammasome complex at the MTOC, as no puncta can be formed without ASC^9,23^. Taken together, our findings here provide a mechanistic basis for future exploration of NLRP3 inflammasome trafficking and activation, as well as inform the development of new anti-inflammatory and anti-cancer treatments targeting this early stage of NEK7-dependent NLRP3 activation.

**Extended Data Fig. 1.**
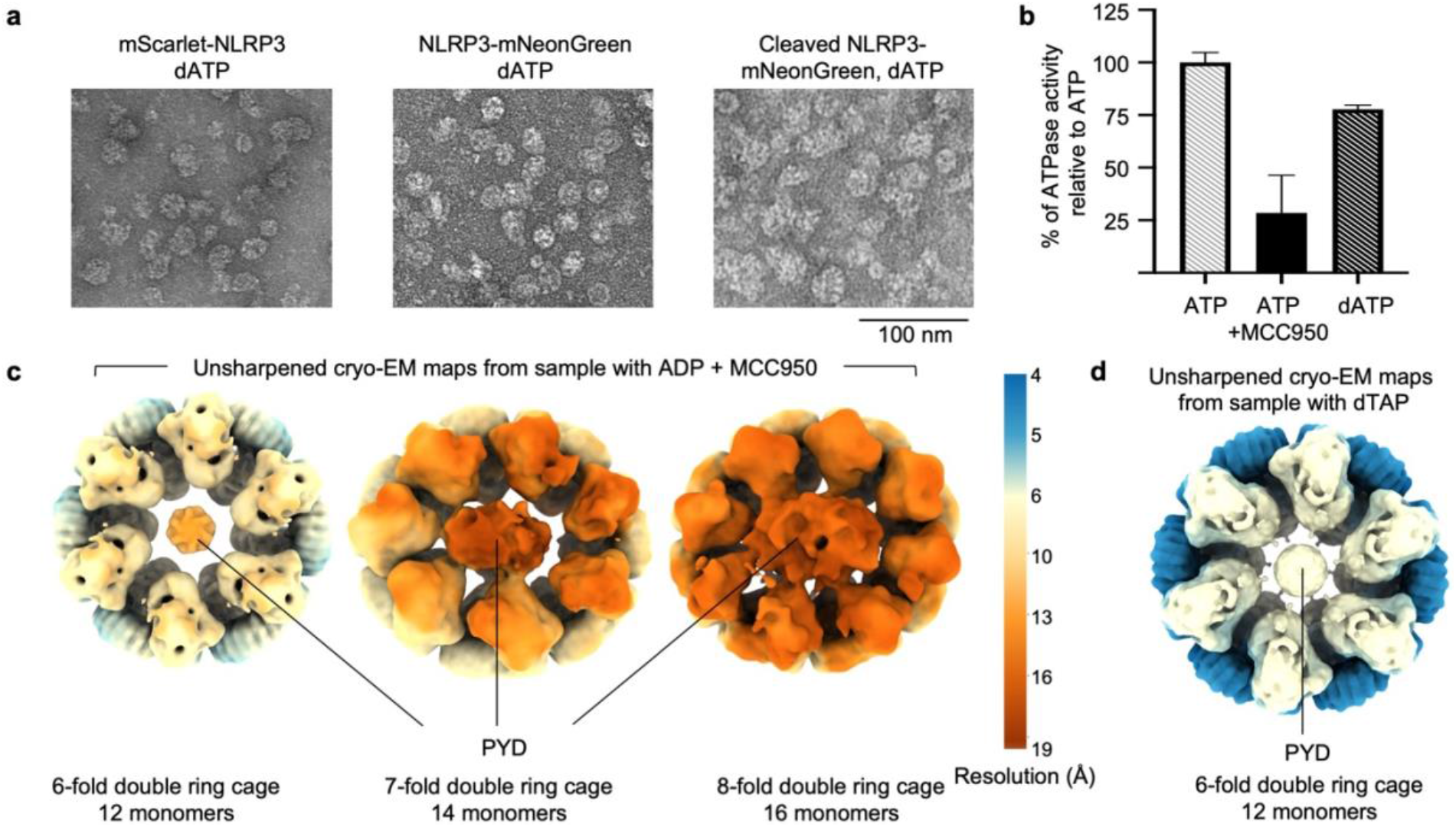
Negative-staining EM imaging and cryo-EM maps of NLRP3 double ring cage. **a**, Representative negative-staining electron microscopy images of NLRP3 oligomers obtained from mScarlet-NLRP3, NLRP3-mNeonGreen and cleaved NLRP3-mNeonGreen constructs. NLRP3 was purified with dATP. **b**, ATPase activity of NLRP3 with ATP, ATP + MCC950 and dATP measured as a percentage of NLRP3 activity with ATP. The assay measured the free phosphate released from the ATPase activity. Data are presented as mean ± s.d., n=3. **c, d**, Unsharpened cryo-EM maps of NLRP3 cage species purified in the presence of ADP and MCC950 (**c**) or dATP (**d**). Maps are colored by resolution. Only the 6-fold double ring cage maps were averaged by D6 symmetry.

**Extended Data Fig. 2.**
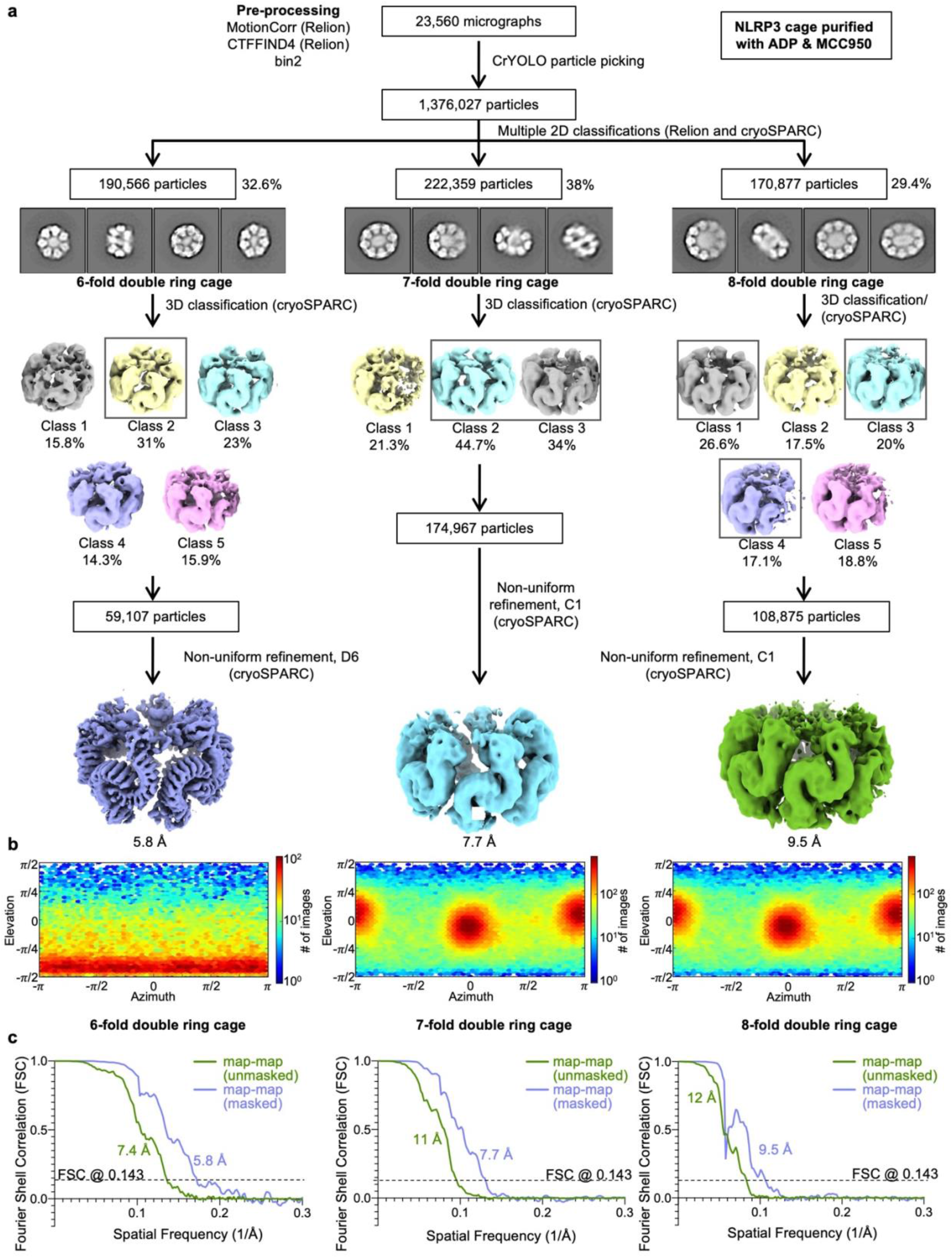
Structure determination workflow for NLRP3 double ring cage purified in the presence of ADP and MCC950. **a**, Workflow for the structure determination with final cryo-EM maps. **b**, Angular distributions for particle projections (cryoSPARC), shown as heat maps of the number of particles for each viewing angle (less=blue, more=red). **c**, Unmasked and masked map-map FSC curves for each cryo-EM map.

**Extended Data Fig. 3.**
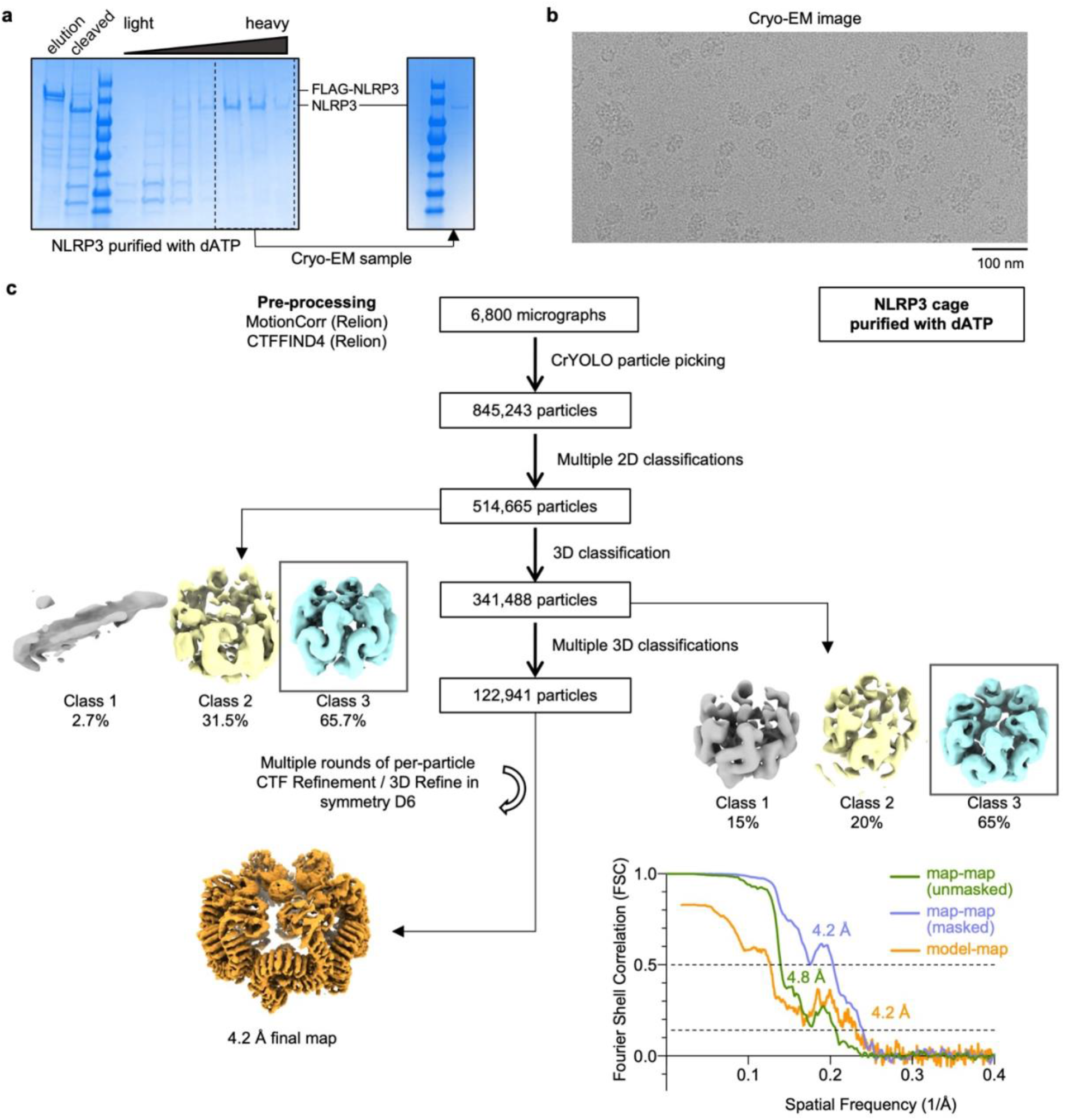
Structure determination workflow for NLRP3 double ring cage purified in the presence of dATP. **a**, SDS-PAGE gel with elution and sucrose gradient fractions of NLRP3 purified from reconstituted HEK293T cells in the presence of dATP (left), fractions collected for the cryo-EM sample (dashed box) and an SDS-PAGE gel of the final sample (right). **b**, A representative cryo-EM micrograph. **c**, Workflow for the structure determination with the final cryo-EM map (bottom left) and the unmasked and masked map-map and model-map FSC curves (bottom right).

**Extended Data Fig. 4.**
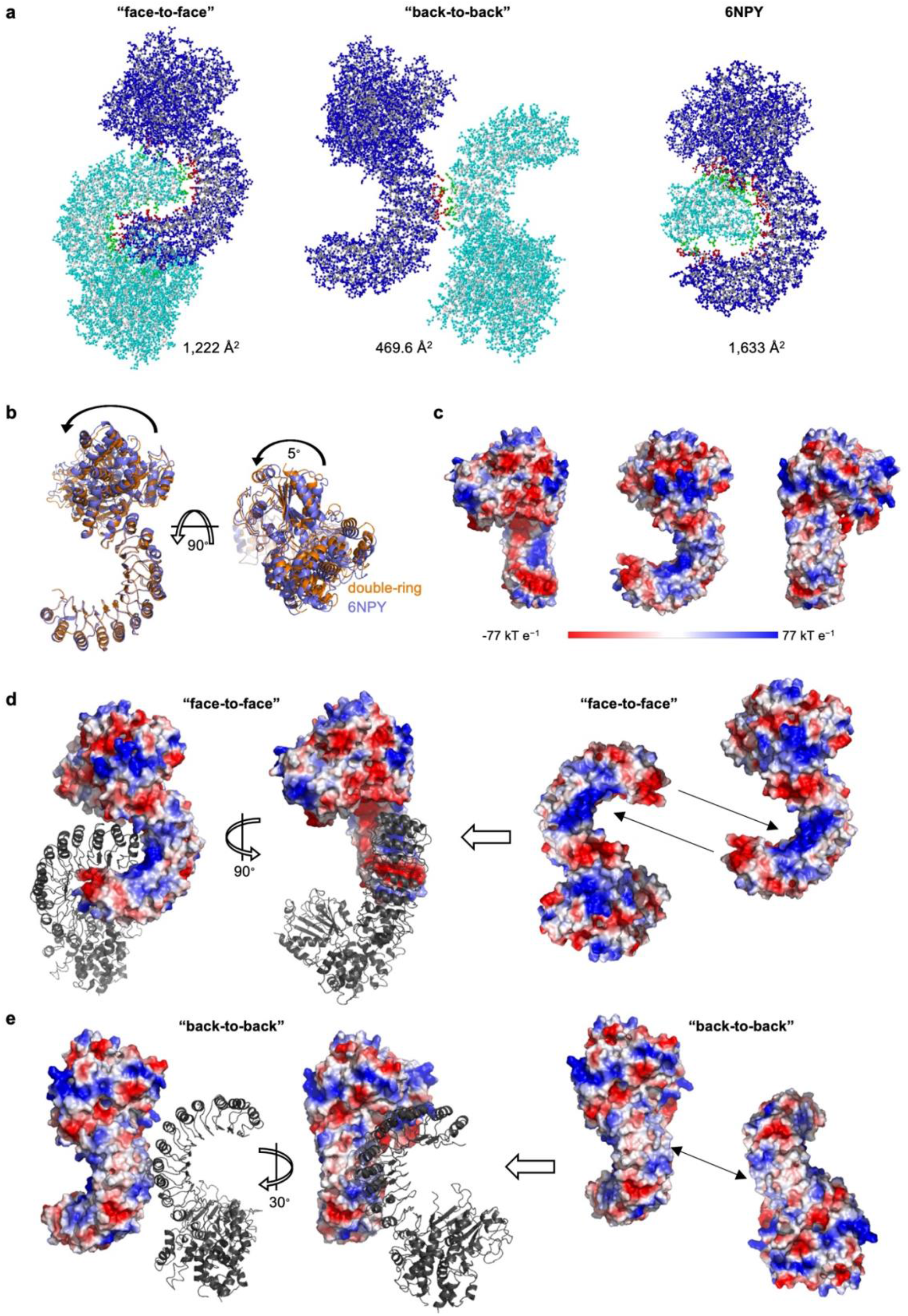
Analysis of interacting interfaces within the NLRP3 double ring cage. **a**, Ball-and-stick models of the interacting monomers within an NLRP3 cage (left and center) and of the NLRP3−NEK7 complex^13^ (6NPY, right). Interface regions are defined by PISA^27^ and colored red and green. **b**, A superposition of an NLRP3 monomer from the double ring cage (orange) with NLRP3 from the NLRP3−NEK7 complex (6NPY, blue). NACHT domain rotation is indicated with an arrow. **c**, Surface electrostatic potential of an NLRP3 monomer from the double ring cage in three orientations, shown in the range of red (−77 kT/e, negatively charged) to blue (77 kT/e, positively charged). **d, e**, “Face-to-face” (**d**) and “back-to-back” (**e**) interfaces. One monomer is colored by electrostatic potential, and the second monomer is colored grey. The “face-to-face” interface is dictated by charge complementarity as shown with arrows. In contrast, the “back-to-back” interface is mostly formed by hydrophobic interactions as shown with arrows.

**Extended Data Fig. 5.**
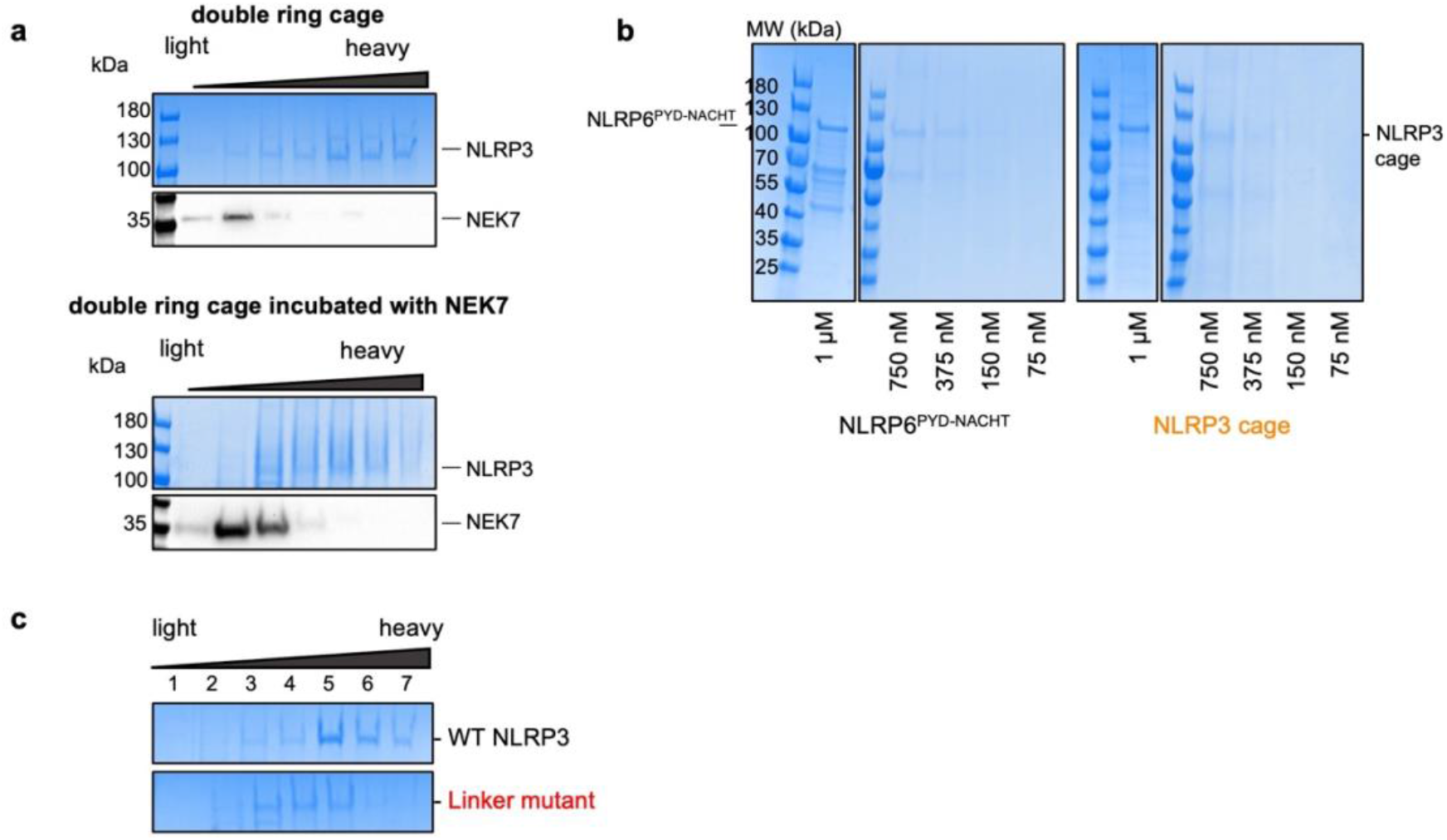
| NEK7 and mutation of the polybasic region disrupt NLRP3 double ring cage formation *in vitro*. **a**, SDS-PAGE of sucrose gradient fractions containing NLRP3 double ring cage purified from reconstituted HEK293T cells in the presence of dATP (“double ring cage”) or NLRP3 cage incubated with excess of recombinant NEK7. NEK7 was visualized with WB using anti-NEK7 antibodies. **b**, An SDS-PAGE gel of NLRP6^PYD-NACHT^ and NLRP3 cage dilutions used for the experiment in Fig. 3a, b. c, SDS-PAGE gel of sucrose gradient fractions containing WT (black), or linker mutant NLRP3 (red). NLRP3 was purified from reconstituted HEK293T in the presence of dATP.

**Extended Data Fig. 6.**
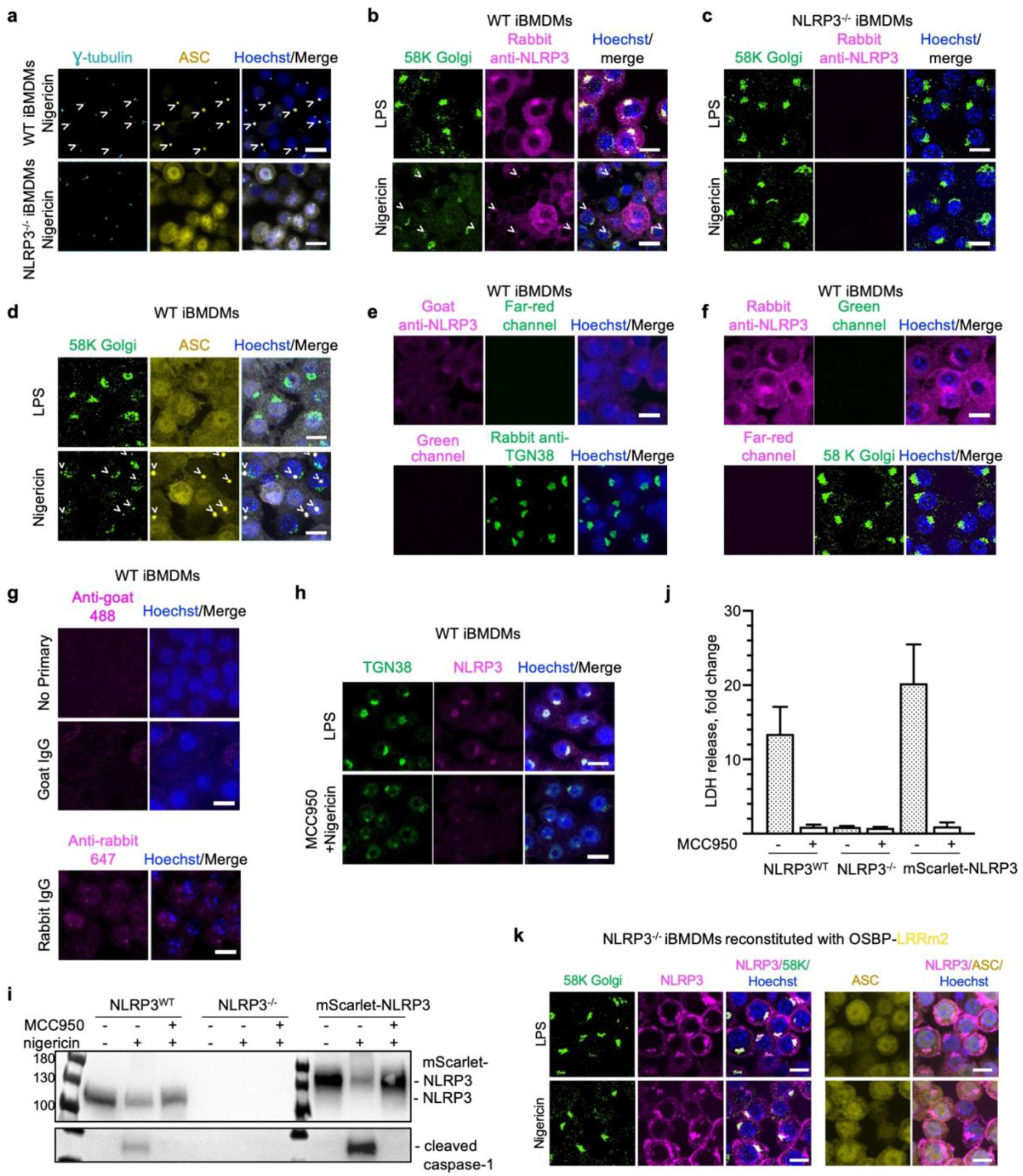
Nigericin induces TGN dispersion in a NLRP3 double ring cage-dependent manner. **a**, Confocal imaging of WT (top) and NLRP3^-/-^ iBMDMs (bottom) primed with LPS and also treated with 20 μM nigericin for 1 h for γ-tubulin (IF, cyan), ASC (IF, yellow), and DNA (Hoechst dye, blue). NLRP3 inflammasome specks are labelled with arrowheads. **b, c**, Confocal imaging of WT (**b**) and NLRP3^-/-^ iBMDMs (**c**) primed with LPS (top) or also treated with 20 μM nigericin for 1 h (bottom) by IF for NLRP3 (rabbit anti-NLRP3 antibody, magenta), 58K Golgi protein (green) and DNA (Hoechst dye, blue). NLRP3 inflammasome specks are labelled with arrowheads. **d**, Confocal imaging of WT iBMDMs primed with LPS (top) or also treated with 20 μM nigericin for 1 h (bottom) by IF for ASC (yellow), 58K Golgi protein (green) and DNA (Hoechst dye, blue). NLRP3 inflammasome specks are labelled with arrowheads. **e, f**, Channel controls for immunofluorescence in (**b-c**) and Fig. 4. WT iBMDMs were visualized by IF for NLRP3 (goat or rabbit anti-NLRP3 antibody, magenta), TGN38 (rabbit anti-TGN38 antibody, green) or 58K Golgi protein (green) in LPS-primed cells. NLRP3 was detected by IF with Alexa488-labeled secondary antibody, TGN38 by IF with Alexa647-labeled secondary antibody, and DNA by Hoechst dye (**e**). NLRP3 was detected by IF with Alexa647-labeled secondary antibody, the 58K Golgi protein by IF with Alexa488-labeled secondary antibody, and DNA by Hoechst dye (**f**). **g**, Isotype controls for the immunofluorescence. LPS-primed WT iBMDMs were incubated with goat IgG, rabbit IgG or no primary antibody and visualized by IF with anti-goat Alexa488- or anti-rabbit Alexa647-labeled antibodies, respectively. **h**, Confocal imaging of WT iBMDMs primed with LPS (top) or also pre-treated with MCC950 and treated with 20 μM nigericin for 1 h (bottom) for TGN38 (IF, green), NLRP3 (mScarlet, magenta) and DNA (Hoechst dye, blue). Z projections with scale bars of 10 μm. **i**, WBs of the whole cell lysates from LPS-, MCC950+nigericin- and nigericin-treated WT iBMDMs, NLRP3^-/-^ iBMDMs, or NLRP3^-/-^ reconstituted with human WT mScarlet-NLRP3. FLAG-mScarlet-NLRP3, NLRP3 and cleaved caspase-1 were visualized with corresponding antibodies. MCC950 inhibited caspase-1 processing. **j**, Cell death indicated by LDH release for WT iBMDMs, NLRP3^-/-^ iBMDMs, and NLRP3^-/-^ iBMDMs reconstituted with WT mScarlet-NLRP3 treated with nigericin only or pretreated with MCC950. The level of LDH release is shown as a fold change between LPS-primed cells treated or not with nigericin. Data are presented as mean ± s.d., n=3. MCC950 abolished NLRP3-mediated cell death. **k**, Confocal imaging of NLRP3^-/-^ iBMDMs reconstituted with OSBP-fused double ring cage disrupting mutant LRRm2 of human mScarlet-NLRP3 primed with LPS (top) or also treated with 20 μM nigericin for 1 h (bottom) for 58K Golgi protein (IF, green), NLRP3 (mScarlet, magenta), ASC (yellow), and DNA (Hoechst dye, blue). Disruption of the double ring cage abolished TGN dispersion upon nigericin treatment even when this NLRP3 mutant was targeted to TGN by the OSBP domain. All images are maximum intensity Z projections with scale bars of 10 μm.

**Extended Data Fig. 7.**
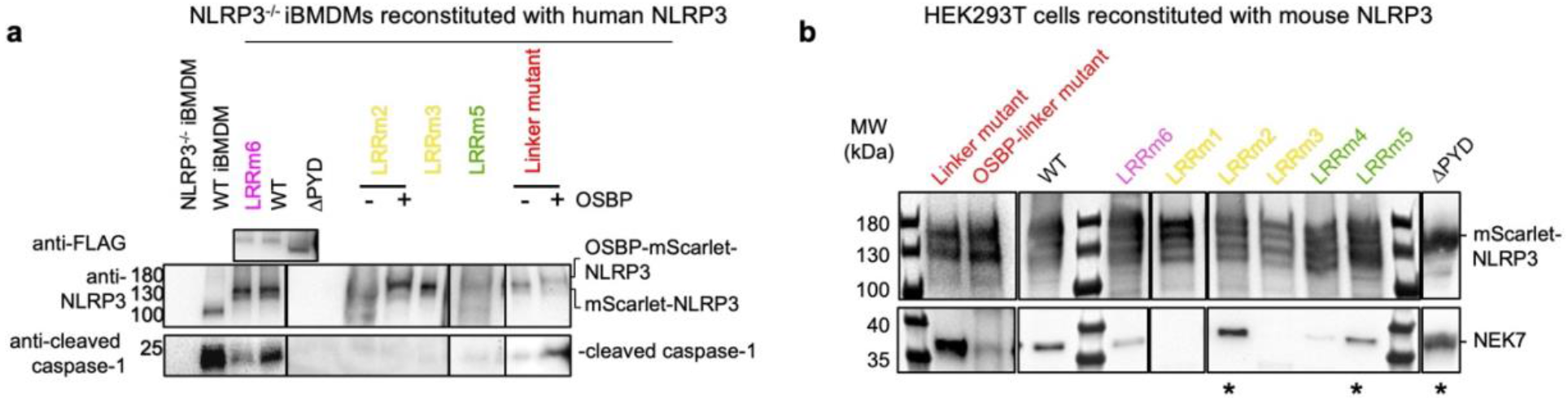
Disruption of NLRP3 double ring cage impairs NLRP3 activity. **a**, WBs of the whole cell lysates from LPS- and nigericin-treated WT iBMDMs, NLRP3^-/-^ iBMDMs, or NLRP3^-/-^ reconstituted with human mScarlet-NLRP3 WT and double ring cage disrupting mutants color coded as in Fig. 2b, c. NLRP3, FLAG-mScarlet-NLRP3 and cleaved caspase-1 were visualized with corresponding antibodies. Double ring cage disruption mutants resulted in reduced caspase-1 processing. **b**, The FLAG pull-down from HEK239T cells reconstituted with mouse FLAG-mScarlet-NLRP3 WT and double ring cage disrupting mutants analyzed by WB using anti-FLAG and anti-NEK7 antibodies. LRRm2, LRRm4, LRRm5 and ΔPYD mutants (asterisks) and linker mutants retained NEK7 biding.

**Extended Data Fig. 8.**
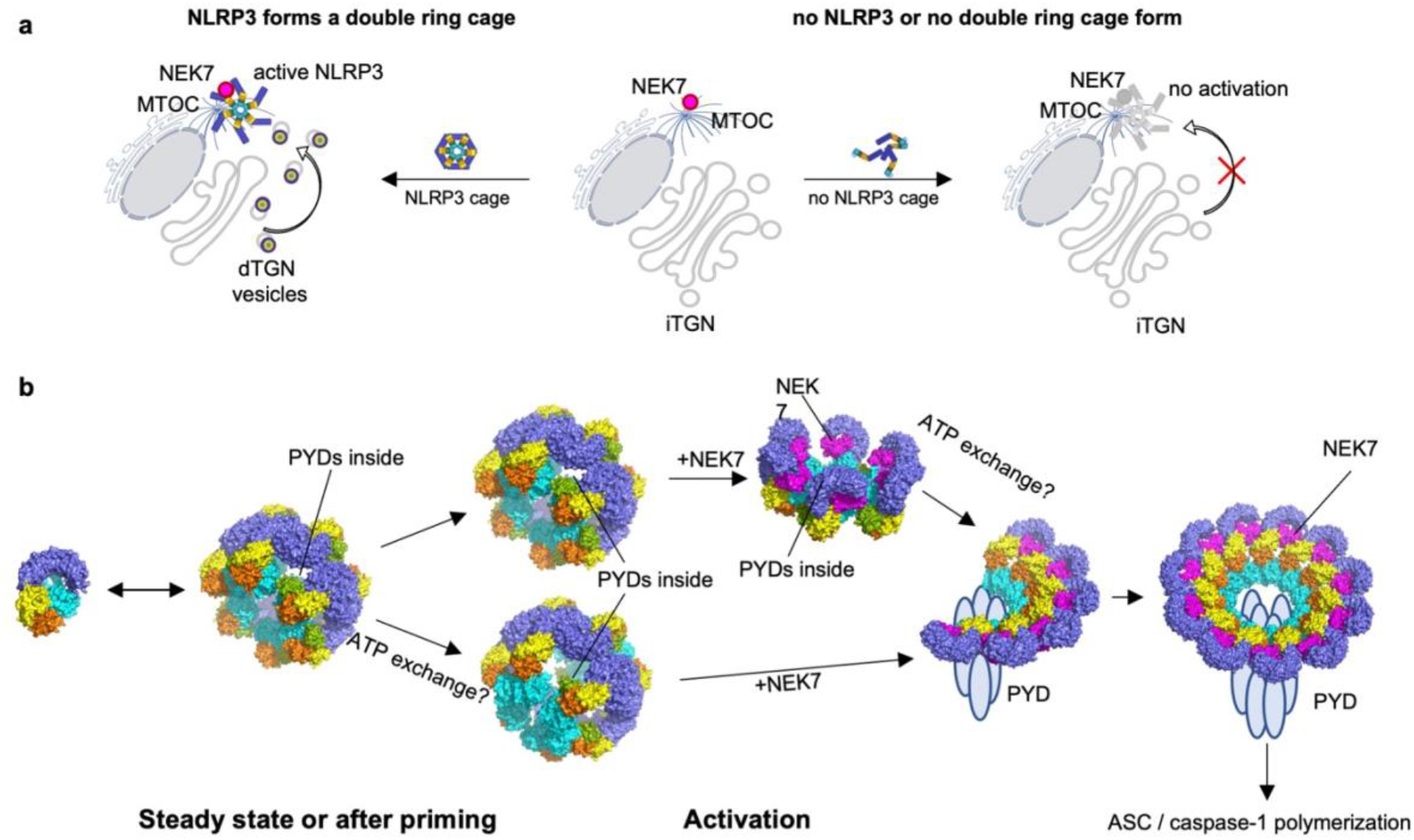
A model for NLRP3 trafficking and activation. **a**, A schematic diagram illustrating the role of the double ring cage in NLRP3 activation. Nigericin or other NLRP3 activating stimuli lead to double ring cage dependent TGN dispersion and transport of TGN vesicles to the MTOC for the association of NLRP3 with NEK7, ASC and caspase-1 (left side). iTGN: intact TGN; dTGN: dispersed TGN. Without double ring cage formation, NLRP3 activating stimuli do not induce TGN dispersion, thus curtailing transport of TGN vesicles to the MTOC and resulting in no NLRP3 activation (right side). **b**, Proposed structural rearrangements of NLRP3 in the course of NLRP3 activation. In a resting state NLRP3 exists presumably in both monomeric and double ring cage forms. Upon activation NLRP3 gets transported to MTOC, with either “closed” (top) or “open” (bottom) NACHT domains. At MTOC the double ring cages get disrupted by NEK7 leading to a partial and then full NLRP3 oligomerization in a form of an active inflammasome complex.

**Extended Data Table 1.**
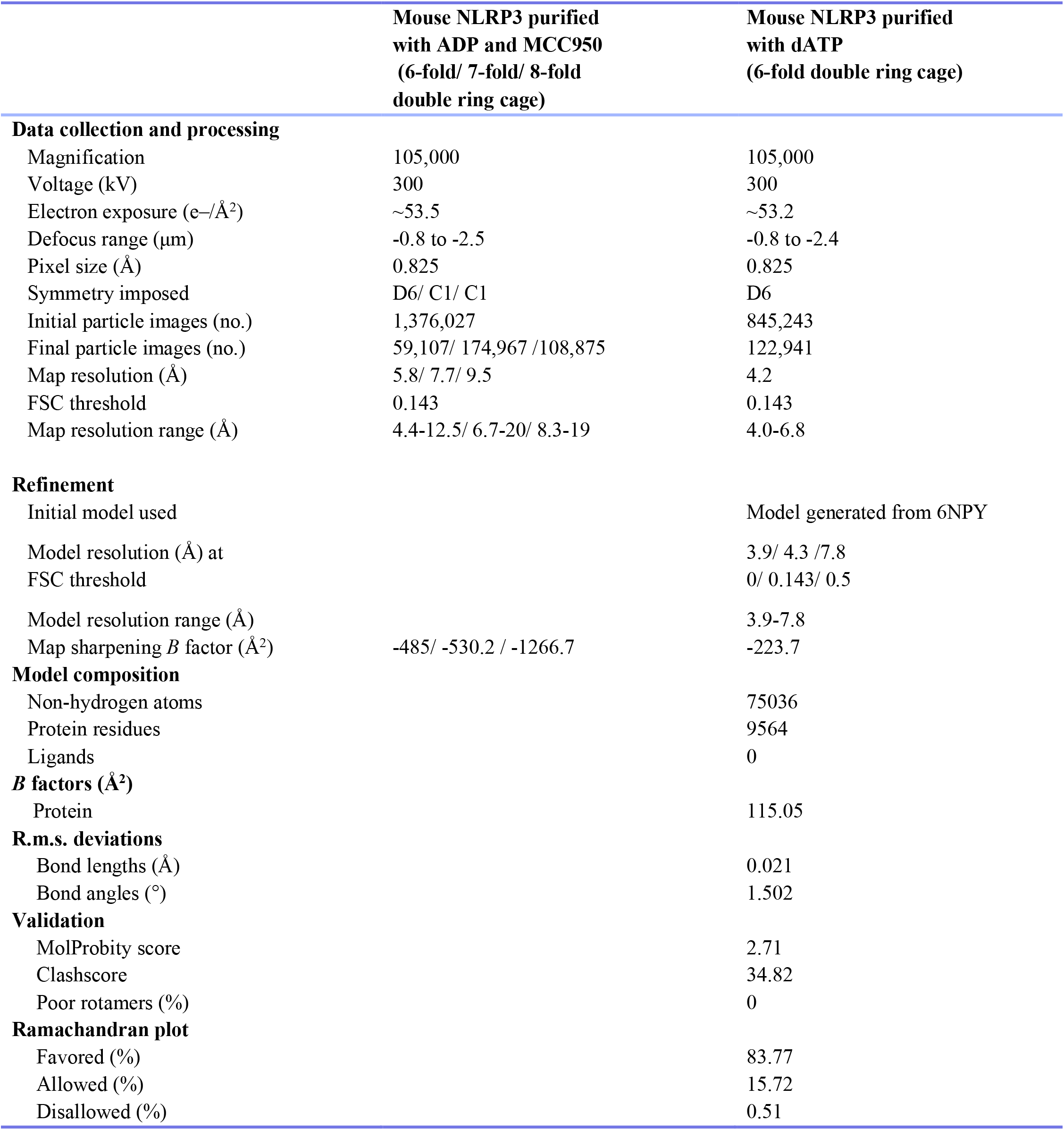
Statistics of NLRP3 double ring cage structure determination by cryo-EM.

## Acknowledgements

We thank Dr. Maria Ericsson at the HMS EM facility for training and support, Drs. Sarah Sterling and Richard Walsh at the Harvard Cryo-EM Center for Structural Biology for cryo-EM training, help with data collection and valuable advice, Dr. Janette Myers at the Pacific Northwest Center for Cryo-EM at Oregon Health & Science University, Dr. Kyounghwan Lee at the University of Massachusetts Cryo-EM Core for screening and preliminary dataset collection, and SBGrid team for software support and computational resources. We also thank Dr. Ross Tomaino and Taplin Biological Mass Spectrometry Facility for protein analysis, Grigoriy Losyev at the BWH Human Immunology Center Flow Core and Ronald Mathieu at ONC-HSCI Flow Cytometry Research Facility for cell sorting, Harry Leung at BCH/PCMM Microscopy Core for training and advice and Paula Montero Llopis at the Microscopy Resources on the North Quad (MicRoN) core at Harvard Medical School for microscope use. WT and NLRP3^-/-^ iBMDMs were kind gifts from Dr. Katherine Fitzgerald at University of Massachusetts Medical School, plasmid pLenti-CMVie-IRES-BlastR (Addgene plasmid #119863) was a gift from Dr. Ghassan Mouneimne, pLV-eGFP (Addgene plasmid #36083) was a gift from Dr. Pantelis Tsoulfas, lentivirus packaging plasmids pMD2.G (Addgene plasmid #12259) and psPAX2 (Addgene plasmid # 12260) were gifts from Dr. Didier Trono. This work was supported by National Institutes of Health (HD087988 and AI124491 to H.W.), the Cancer Research Institute Irvington Postdoctoral Fellowship (to C.S.) and Damon Runyon Fellowship Award (to L.A.).

## Author contributions

L.A and H.W. designed and conceptualized the study. P.P. provided initial NLRP3-expressing HEK293 cells. L.A. preformed cloning, protein purifications, negative staining EM, cryo-EM sample preparation, screening and data collection, generation of stable cell lines, cytotoxicity assays and immunoblotting, fractionation studies and biochemical evaluation of the NLRP3 complex in sucrose gradients, ASC^PYD^ polymerization assay and native-PAGE. L.D. performed immunofluorescence microscopy experiments, image analysis and quantifications of microscopy data. S.R. and L.A. analyzed cryo-EM data and performed model building and refinement. C.S. assisted with protein purification and ASC^PYD^ polymerization assay. T.P. performed ATPase activity assay and assisted with protein purification with L.A.’s input. L.A. and H.W. wrote the manuscript with input from all authors.

## Methods

### Constructs and cloning

Full-length mouse and human NLRP3 were cloned into pLenti CMVie-IRES-BlastR (Addgene plasmid #119863). The resulting constructs were further modified by addition of N-terminal FLAG-tag followed by fluorescent protein mScarlet and Tobacco Etch Virus (TEV) protease cleavage site upstream of the NLRP3 gene. An additional N-terminal FLAG and pleckstrin homology domain of oxysterol-binding protein 1 (OSBP) domain (aa 87–185) were added to selected constructs. Full-length mouse NLRP3 was also amplified with addition of N-terminal FLAG and C-terminal TEV sequences and cloned into an in-house modified pLV-eGFP vector, in which eGFP was replaced by mNeonGreen (mNG), for FLAG-NLRP3-TEV-mNG expression. Full length mouse NEK7 were cloned into a modified pET28a vector with an N-terminal 6×His-SUMO tag followed by the Ulp1 protease site. Constructs encoding monomeric MBP-tagged NLRP3^NACHT-LRR^ (aa 134-1034) and containing human NLRP6^PYD-NACHT^ (aa 1-564) in pDB-His-MBP vector were described before^13,32^. Monomeric MBP-tagged NLRP3^NACHT-LRR^ (aa 134-1034) was subcloned into pLenti CMVie-IRES-BlastR (Addgene plasmid #119863) with addition of N-terminal FLAG-tag. The S106C mutant of human ASC PYD (aa1-106) was cloned into an in-house modified pET28-MBP-TEV vector (Addgene, plasmid #69929), in which N-terminal His6 tag was added for expression of proteins with a cleavable N-terminal His_6_-MBP-tag, based on previously described constructs^35^. The ASC^PYD^ S106C mutant was used for thiol-labelling by Alexa488. Point mutations were introduced with a QuikChange site-directed mutagenesis protocol using Q5 High-Fidelity 2x Master Mix (NEB). NLRP3 mutants were named: LRRm1 for mouse Y858C_W830A and human W833A_Y861C, LRRm2 for mouse N1008R_R1009E_E1010R_R1013E and human N1011R_Y1012A_R1013E_S1016A, LRRm3 for mouse R771E_W773T_R776E and human R774E_W776A_R779E, LRRm4 for mouse D809R_F810A_R813E and human D812R_F813A_R816E, LRRm5 for mouse H781E_Q782R_F785A and human H784E_E785R_F788A, and LRRm6 for mouse K970E_ Q1001R_ F1029A and human K973E_F1032A.

### Cell culture and generation of stable cell lines

WT and NLRP3-knockout (NLRP3^-/-^) immortalized bone marrow–derived macrophages (iBMDMs) were a gift from Prof. K. A. Fitzgerald (University of Massachusetts Medical School, Worcester, MA, USA). HEK293T cells and iBMDMs were cultivated in Dulbecco’s Modified Eagle’s medium (DMEM), with L-glutamine (Thermo Fisher Scientific, Cat. no: 10569-004), supplemented with 10% fetal bovine serum (Thermo Fisher Scientific, Cat. no: 16000-044) at 37°C in 5% CO_2_. Cells reconstituted with pLenti CMVie-IRES-BlastR-based constructs were maintained with addition of 5 µg/ml and 1 µg/ml blasticidin (InvivoGen, Cat. no: ant-bl-05) for HEK293T and iBMDMs, respectively.

For lentivirus preparation HEK293T cells were transfected with 1 µg of plasmid containing the construct of interest, 750 ng psPAX2 packaging plasmid and 250 ng pMD2.G envelope plasmid (Addgene plasmids #12260 and #12259). On the following day the virus-containing medium was collected and filtered using 0.45 µm filter (Millipore Sigma, Cat. no: SLHVM33RS). Fresh HEK293T cells were infected by adding the lentivirus-containing medium to culture medium followed by 1 to 2 day incubation. iBMDMs NLRP3^-/-^ were infected with the spinfection protocol as follows. The cells were resuspended in the lentivirus-containing medium with addition of 8 μg/ml polybrene (Santa Cruz Biotechnology, Cat. no: sc-134220) and centrifuged for 90 min at 2,500 g at room temperature. The supernatant was discarded, and infected cells were incubated in fresh medium for 48 h followed by the second round of spinfection. Positive cells expressing mScarlet-tagged constructs were selected by cell sorting at BD FACSAria Fusion or FACSAria II cell sorter equipped with 100 µm nozzle 20 psi or 85 µm nozzle 45 psi depending on the experiment. The sorted populations were gated to exclude dead and non-fluorescent cells.

### Protein expression and purification

Full-length mouse NEK7 was purified as described before^13^. Full-length mouse NLRP3 was purified from 240 150 mm dishes of HEK293T cells stably expressing FLAG-mScarlet-tagged NLRP3. Trials with HEK293 suspension cells (293F and Expi293) did not yield well behaving NLRP3 proteins. Cells were harvested at 80-90% confluency, washed with phosphate buffered saline (PBS), pelleted, flash-frozen in liquid nitrogen, and stored at -80 °C. Cell pellets were resuspended in a buffer containing 30 mM HEPES, 150 mM NaCl, 5 mM MgCl_2_ and 10% glycerol, pH 7.5, supplemented with protease inhibitor cocktail (Sigma, Cat. no: S8830) and 1 mM tris(2-carboxyethyl)phosphine (TCEP) shortly before usage. Cells were sonicated (3 s on, 8 s off, 3 min total on, 40% power, Branson), centrifuged at 42,000 RPM for 1 h (45 Ti fixed-angle rotor, Beckman), and the supernatant was used for affinity chromatography with anti-FLAG M2 affinity gel (Sigma, Cat. no: A2220) by gravity flow. For purification from the membrane, the pellet after sonication was resuspended in the lysis buffer supplemented with 1% (w/v) n-Dodecyl β-D-maltoside and incubated for 2 h. This step was then followed by centrifugation at 21,000 g for 30 min, and the supernatant was used for affinity purification with anti-FLAG M2 affinity gel (Sigma, Cat. no: A2220). The beads were washed with 50 column volumes (CV) of the same buffer and the protein was eluted with 100 μg/ml 3xFLAG peptide (ApexBio, Cat. no: A6001). The eluted protein was incubated with 0.5 mM ADP and 0.5 mM MCC950 (Millipore Sigma, Cat. no: 538120), 0.5 mM dATP or no additives for 30 min on ice, followed by incubation or not with TEV protease for 30 min at 30 °C to remove the tag. For fig. S10 the elution was additionally incubated with 10 μM NEK7 for 5 h on ice. The mixture was loaded onto a step-gradient of 20%, 25%, 30%, 35%, 40%, 45% and 50% sucrose in 30 mM HEPES at pH 7.5 and 150 mM NaCl, supplemented with protease inhibitor cocktail (Sigma, Cat. no: S8830) and 1 mM TCEP and centrifuged for 16 h at 40,000 rpm (MLS-50 swinging-bucket rotor, Beckman). Fractions of 750 μl were collected manually and NLRP3-containing fractions were merged and buffer-exchanged with Zeba™ spin desalting columns (fisher Scientific, Cat. no: PI87771) equilibrated with the buffer containing 30 mM HEPES at pH 7.5, 150 mM NaCl, 5 mM MgCl_2_, protease inhibitor cocktail, 1 mM TCEP, supplemented with 0.2 mM of the same nucleotides or additives used prior to the sucrose gradient. NEK7 in these fractions was visualized by Western blotting using mouse anti-NEK7 antibody (1:2,000, Santa Cruz, Cat. no: sc-393539) and anti-mouse IgG conjugated to horseradish peroxidase (HRP) (1:2,000, Cell Signaling, Cat. no: 7076S).

For Extended Data Fig. 1 MBP-tagged NLRP3^NACHT-LRR^ was expressed in Sf9 cells using a baculovirus system and purified as described previously^13^. One liter of Sf9 with density of 3×10^6^ cells/ml was infected with 1% v/v of baculovirus. 48 h post-infection cells were collected and lysed by sonication (3 s on, 8 s off, 3 min total on, 40% power, Branson) in a buffer containing 30 mM HEPES at pH 7.5, 200 mM NaCl and 10% glycerol, supplemented with protease inhibitor cocktail (Sigma, Cat. no: S8830) and 1 mM TCEP shortly before usage. The lysate was centrifuged at 42,000 RPM for 1 h (45 Ti fixed-angle rotor, Beckman), and the supernatant was used for affinity chromatography with MBP beads by gravity flow. The protein was eluted with 10 mM maltose solution in the lysis buffer and further purified by size exclusion chromatography using Superose 6 10/300 GL column (GE Healthcare) equilibrated with 30 mM HEPES at pH 7.5, 150 mM NaCl, and 1 mM TCEP.

For Fig. 3 FLAG- and MBP-tagged NLRP3^NACHT-LRR^ was expressed in expi293F cells. One liter of 3×10^6^ cells/ml was transfected with 1 mg plasmid using polyethylenimine (3 mg/l) as transfection reagent. 24 h post-transfection cells were supplemented with glucose (9 mL, 45%) and valproic acid (10 mL, 300 mM) and harvested 4 days after transfection. Cells were lysed by sonication (3 s on, 8 s off, 3 min total on, 40% power, Branson) in a buffer containing 30 mM HEPES, 200 mM NaCl and 10% glycerol, pH 7.5, supplemented with protease inhibitor cocktail (Sigma, Cat. no: S8830) and 1 mM TCEP shortly before usage. The lysate was centrifuged at 42,000 RPM for 1 h (45 Ti fixed-angle rotor, Beckman), and the supernatant was used for affinity chromatography with anti-FLAG M2 affinity gel (Sigma, Cat. no: A2220) by gravity flow. The protein was eluted with 100 μg/ml 3xFLAG peptide (ApexBio, Cat. no: A6001), incubated with an excess of NEK7 for 1h on ice and further purified by size exclusion chromatography using Superose 6 10/300 GL column (GE Healthcare) equilibrated with 30 mM HEPES, 150 mM NaCl, pH 7.5 and 1 mM TCEP.

NLRP6^PYD-NACHT^ protein was expressed and purified as described before^32^. In short, the protein was expressed in *E. coli* BL21 (DE3) at 18 °C overnight following the induction with 0.1 mM isopropyl-β-D-thiogalactopyranoside (IPTG). Cells were lysed by sonication in 20 mM HEPES at pH 7.4, 300 mM NaCl and 20 mM imidazole, centrifuged at 17,000 rpm for 1 h (SA-600, Sorvall) and the supernatant was used for Ni-NTA affinity chromatography. The protein was further purified by size exclusion chromatography using Superdex 200 10/300 GL column (GE Healthcare) equilibrated with 20 mM HEPES at pH 7.4, 150 mM NaCl and 1 mM TCEP.

ASC^PYD^ with the S106C mutation was expressed as a His_6_-MBP fusion protein in *E. coli* BL21 (DE3) at 18 °C overnight following the induction with 0.1 mM IPTG. Cells were lysed by sonication in a buffer containing 30 mM HEPES at pH 7.5, 200 mM NaCl, 10% Glycerol, supplemented with protease inhibitor cocktail and 1 mM TCEP and centrifuged at 40,000 RPM for 1 h (45 Ti fixed-angle rotor, Beckman). Supernatant was incubated with amylose beads pre-equilibrated in the same buffer for 1 h at 4 °C and loaded onto a gravity flow column. The beads were washed with 50 CV of the same buffer and eluted with 50 mM maltose. The eluted protein was fluorescently labelled at the C106 thiol group by incubation with a 2-fold access of AlexaFluor 488 C_5_-maleimide (ThermoFisher, Cat. no: A10254) at 4 °C overnight. The labelled protein was further purified by size-exclusion chromatography using Superdex 200 10/300 GL column (GE Healthcare) equilibrated with 30 mM HEPES at pH 7.5, 150 mM NaCl, supplemented with protease inhibitor cocktail and 1 mM TCEP.

### Negative-staining electron microscopy

For negative staining 5 µl of NLRP3 sample was placed on a copper grid (Electron Microscopy Sciences, cat. no: FCF400CU50), incubated for 1 min, washed twice with buffer containing 30 mM HEPES at pH 7.5 and 150 mM NaCl, stained with 2% uranyl formate for 30 sec and air-dried. The images were collected at a Tecnai G2 Spirit BioTWIN Transmission Electron Microscope (TEM) equipped with AMT 2k CCD camera at 49,000x magnification and 120 keV (HMS EM core facility).

### Cryo-EM data collection

A 4 µl drop containing NLRP3 complex was applied to a Lacey Carbon grid with ultrathin carbon support (Ted Pella, Cat. no: 01824G), incubated for 1-1.5 min, blotted for 3 s, plunged into liquid ethane, and flash frozen using a FEI Vitrobot Mark IV at 100% humidity and 4 °C. Grid conditions were optimized during extensive screening at Pacific Northwest Center for Cryo-EM at Oregon Health & Science University (PNCC), the University of Massachusetts Cryo-EM Core (UMASS) and the Harvard Cryo-EM Center for Structural Biology (HMS) using FEI Talos Arctica (ThermoFisher) microscopes equipped with an autoloader (200 keV, Gatan K3 direct electron detector). Detailed cryo-EM data collection settings are summarized in Extended Data Table 1.

Final datasets were collected at HMS using a Titan Krios microscope (ThermoFisher) operating at 300 keV and equipped with a BioQuantum Imaging Filter (Gatan) and K3 direct electron detector (Gatan). Automated data collection was performed using SerialEM^46^ software, and the movies were obtained in counting mode at 105,000x magnification (0.825 Å/pix). Data quality for each dataset was assessed with on-the-fly data processing algorithm designed by Shaun Rawson for HMS facility. For NLRP3 sample purified in the presence of ADP and MCC950 23,560 movies with 50 frames each were recorded at multiple defocus values from -0.8 to -2.5 μm and with multiple exposures per stage shift (5×4) introduced with image shift. Each movie was acquired at a dose rate 18.194 e/s per physical pixel and accumulated a total dose of 53.46 e/Å^2^ over 2 s total exposure time. For NLRP3 sample purified in the presence of dATP 6,800 movies with 50 frames each were recorded at multiple defocus values from -0.8 to -2.4 μm and with multiple exposures per stage shift (5×4) introduced with image shift. Each movie was acquired at a dose rate 23.951 e/s per physical pixel and accumulated a total dose of 53.225 e/Å^2^ over 1.51 s total exposure time.

### Cryo-EM data processing

Cryo-EM data processing software and support was provided by SBGrid consortium^47^. Raw movies were corrected by gain reference and beam-induced motion and combined into a motion-corrected micrograph using the MotionCorr2 algorithm^48^. The defocus value for each micrograph was determined with CTFFIND4^49^. Automated particle picking was performed with crYOLO^50^ using a general model (optimized in HMS facility).

A dataset of NLRP3 purified in the presence of ADP and MCC950 yielded 1,376,027 particles picked by crYOLO^50^, which were extracted with 2x binning resulting in 1.65 Å pixel size. Initial rounds of 2D classifications were performed in Relion3.1^51,52^, after which the 2D classes were manually sorted into 6-fold, 7-fold and 8-fold species (190,566, 222,359 and 170,877 particles, respectively), imported into cryoSPARC^53^ and processed separately. For each species 3 initial models were generated with *ab initio* reconstruction. The best model was then used for 3D classification after a 35 Å low-pass-filtering in cryoSPARC^53^ using the heterogenous refinement mode for 3 or more outputs. The best 3D classes with the least particle distortion were merged and used for the non-uniform 3D refinement using symmetry D6 for a 6-fold double ring cage or C1 for 7- and 8-fold double ring cages, which resulted in final cryo-EM maps at 5.8 Å, 7.7 Å and 9.5 Å resolutions generated from 59,107, 174,967 and 108,875 particles, respectively. The described cryo-EM workflow for this dataset is also presented in Extended Data Fig. 2 with gold-standard Fourier shell correlation (FSC) for each map. Local resolution estimation was calculated with Local Resolution Estimation module in cryoSPARC^53^. Post-processing of a 6-fold double ring cage map was performed with DeepEMhancer^54^.

A dataset of NLRP3 purified in the presence of dATP yielded 845,243 particles picked by crYOLO^50^, which were extracted with no binning resulting in 0.825 Å pixel size. Further processing was performed in Relion 3.1^51,52^. The extracted particles were subjected to multiple 2D classifications until the classes appeared homogenous. Initial model was built with *de novo* reconstruction, low-pass-filtered to 40 Å and used for subsequent multiple rounds of 3D classification and refinement. The refinement of the best class containing 122,941 particles using D6 symmetry resulted in a 4.3 Å map, which could be further improved up to 4.2 Å by multiple rounds of per-particle CTF refinement followed by 3D refinement in D6. Bayesian polishing or beam tilt CTF refinement did not result in any further improvement^55,56^. The final map was post-processed in Relion 3.1^51,52^ for calculation of FSC curves and DeepEMhancer^54^ for model building and representation. The described cryo-EM workflow for this dataset is also presented in Extended Data Fig. 3. Local resolution estimation was calculated with Relion 3.1^51,52^.

### Model building and structure representation

The atomic model was built using cryo-EM map obtained from the NLRP3 complex purified in the presence of dATP (4.2 Å). 12 molecules of NLRP3^NACHT-LRR^ without NEK7 from the structure of the NLRP3:NEK7 complex^13^ (PDB ID 6NPY) were first fit into the cryo-EM map using UCSF-Chimera^57^ rigid body fit. Each chain was then split into LRR (aa 694-1034) and NACHT (aa 133-693) segments, which were fitted separately into the cryo-EM map in UCSF-Chimera^57^. Residues corresponding to the mouse NLRP3 sequence different from the human model were mutated in Coot^58^ and the rotamers were selected to position the side chains close to those from 6NPY NLRP3 model. Due to the modest resolution (4.2 Å) no further manual adjustments of the model were performed and the model was subjected to real-space refinement in Phenix^59,60^ with the starting model as a reference. The final model represents a mouse NLRP3 dodecamer and includes residues Asp131-Trp1033 of each monomer. Interface analysis of the resulting model was performed with PISA^27^. Structure representations were generated using ChimeraX^61^ and Pymol^62^.

### ATPase assay

10 µM NLRP3^NACHT-LRR^ were mixed with 5 mM ATP or dATP, or 5 mM ATP and 100 µM MCC950 in a buffer containing 30 mM HEPES at pH 7.5, 150 mM NaCl and 15 mM MgCl_2_, and incubated at room temperature. At 0 min and 1 h the sample was analyzed for concentrations of inorganic monophosphates with the ATPase/GTPase Activity Kit (Sigma-Aldrich, Cat. no: MAK113-1KT) according to manufacturer’s instructions.

### ASC^PYD^ polymerization assay by fluorescence quenching

2 µM of Alexa488-conjugated His_6_-MBP-TEV-ASC^PYD^ (S106C) was mixed with NLRP3 cage or NLRP6^PYD-NACHT^ at different concentrations in buffer containing 30 mM HEPES at pH7.5, 150 mM NaCl, 5 mM MgCl_2_, 0.05% Triton X100 and 0.2 mM dATP, in black round-bottom 384-well plate (Corning, Cat. no: 3820) with a 20 µl final volume. Reaction was initiated by addition of TEV protease at 0.5 mg/ml final concentration. Fluorescence was measured at 25 °C using Synergy™ NEO HTS Multi-Mode Microplate Reader (BioTek) with excitation and emission wavelengths 420 nm and 485 nm, respectively. The initial linear intervals of the fluorescence curves were used to calculate the polymerization rate for each sample as relative fluorescence units (RFU)/minute.

### *In vitro* lipid blot assay

Lipid binding assay with purified NLRP3 variants was performed using PIP strip (Echelon Biosciences, Cat. no: P-6001) according to manufacturer instructions. In short, the PIP strip membranes were blocked using 3% bovine serum albumin (BSA) in phosphate-buffered saline with 0.1% Tween 20 (PBS-T) for 1 h following by incubation with 30 µl NLRP3 FLAG-tagged proteins (0.5 µM) diluted in 3% BSA in PBS-T for 1 h. Next, the membranes were washed in PBS-T for 40 min and the bound proteins were visualized with anti-FLAG-HRP antibodies (1:10000 in PBS-T for 1h, Sigma-Aldrich, Cat. no: A8592). All steps were performed at room temperature. All samples were analysed at the same time under the same conditions.

### Native PAGE

WT and NLRP3^-/-^ iBMDMs were seeded on 150 mm plates at 5×10^6^ cells/plate and on the next day treated with 1 μg/ml LPS (Invivogen, Cat. no: tlrl-b5lps) for 3.5 h.. Untreated and LPS-treated cells were collected, washed with PBS and resuspended in a buffer containing 30 mM HEPES at pH 7.5, 150 mM NaCl, 5 mM MgCl_2_ and 10% glycerol, followed by sonification (3 s on, 8 s off, 1 min total on, 40% power, Branson) and centrifugation at 36,000 rpm (MLA-50 fixed-angle rotor, Beckman) for 1 h. Membrane pellet was resuspended in the same buffer and protein extraction was performed by adding 1% n-Dodecyl β-D-maltoside (w/v) followed by incubation for 2 h and centrifugation at 21,000 g for 30 min. Supernatants were loaded on a 3-12% Bis-Tris gel (ThermoFisher, Cat. no: BN1003BOX) and the Native PAGE was run at 100 V for 4 h at 4 °C. NLRP3 and β-actin were visualized by Western blotting using anti-NLRP3 (1:2,000, Adipogen, Cat. no: AG-20B-0014-C100) and anti-β-actin (1:2,000, Sigma Aldrich, Cat. no: A2228-100UL) primary antibodies, respectively, followed by anti-mouse-HRP (1:5,000, Cell Signaling, Cat. no: 7076S) secondary antibody.

### Immunoblotting of whole cell lysates

WT iBMDMs, NLRP3^-/-^ iBMDMs or NLRP3^-/-^ iBMDMs reconstituted with mScarlet-tagged NLRP3 were seeded at 0.5×10^6^ cells/well on a 12-well tissue culture plate and on the next day treated or not with 1 μg/ml LPS (Invivogen, Cat. no: tlrl-b5lps) for 3.5 h or with 1 μg/ml LPS for 2.5 h and additionally with 20 μM MCC950 (Millipore Sigma, Cat. no: 538120) for 1h followed by NLRP3 activation with 20 μM nigericin (Sigma-Aldrich, Cat. no: N7143-5MG) for 1 h. The medium was discarded and the whole cell lysates were prepared by adding SDS sample loading buffer (50 mM Tris at pH 7.5, 2% w/v SDS, 10% glycerol, 150 mM β-mercaptoethanol, 1.5 mM bromophenol Blue) directly to the cells. NLRP3, β-actin and cleaved caspase-1 p20 fragment were visualized by Western blotting using anti-NLRP3 (1:2000, Adipogen, Cat. no: AG-20B-0014-C100), anti-β-actin (1:2,000, Sigma Aldrich, Cat. no: A2228-100UL) and anti-p20 of caspase-1 (1:1,000, Cell Signaling, Cat. no: 89332S) primary antibodies, respectively. Anti-mouse-HRP (1:5000, Cell Signaling, Cat. no: 7076S) and anti-rabbit-HRP (1:2,500, Cell signaling, Cat. no: 7074S) secondary antibodies were used.

### Immunofluorescence (IF)

Cell lines were plated on CELLview 4-compartment dishes (Greiner Bio-One), treated with 1 μg/ml LPS (Invivogen, Cat. no: tlrl-b5lps) for 3.5 h followed or not by NLRP3 activation with 20 μM nigericin (Sigma-Aldrich, Cat. no: N7143-5MG) for 1h with or without a pretreatment with 20 μM MCC950 (Millipore Sigma, Cat. no: 538120) for 1h prior to activation, fixed in 3% paraformaldehyde (PFA) for 30 min at 4 °C and permeabilized with 0.1% Triton X-100 for 10 minutes. Cells were incubated in PBS-Tween containing 3% BSA for 3 h, which minimized non-specific binding. After three washes with PBS-Tween, cells were incubated overnight at 4 °C with primary antibodies. The following antibodies were used: rabbit monoclonal anti-NLRP3 (1:250, Abcam, Cat. no: ab272702), goat polyclonal anti-NLRP3 (1:100, Abcam, Cat. no: 4207), rabbit polyclonal anti-TGN38 (1:500, Novus Biologicals, Cat. no: NBP1-03495), rabbit polyclonal anti-TGN46 (1:500, Abcam, Cat. no: 50595), mouse monoclonal anti-58K Golgi (1:500, Novus Biologicals, Cat. no: NB600-412), sheep polyclonal anti-TGN38 (1:500, Novus Biologicals, Cat. no: NBP1-20263), rabbit polyclonal anti-ASC (1:1000, Cell Signaling, Cat. no: 67824) and mouse monoclonal anti-γ-tubulin (1:500, Abcam, Cat. no: ab11316). After incubation, cells were washed and incubated with AlexaFluor647-conjugated anti-rabbit IgG (1:500, ThermoFisher, Cat. no: A27040) and/or Alexa488-conjugated anti-mouse IgG (1:500, ThermoFisher, Cat. no: ab150113) for 1 h at room temperature, washed with PBS (3 × 10 min) and then stained with Hoechst (1:500, Immunochemistry Technologies, Cat. no: 639). For negative controls we used rabbit IgG (Santa Cruz, Cat. no: 2027), goat IgG (Santa Cruz, Cat. no: 2028) and sheep IgG (R&D, Cat. no: 5-001-A) at the same concentration for primary antibodies. Moreover, we performed staining with no primary antibodies as an additional control. Cells were imaged using a Nikon Ti inverted microscope fitted with a Photometrics CoolSNAP HQ2 Peltier cooled CCD camera and Andor Zyla 4.2 sCMOS camera equipped with Plan Apo 100x/1.4 Ph3, Plan Apo 60x/1.3 DIC and Plan Apo 40x/1.3 DIC objectives. Lumencor SpectraX LED illuminator was used. Chroma 49000 (DAPI), Chroma 29002 (green), Chroma 49008 (red) and Chroma 49011 (far red) filter cubes were used. Image analysis was performed in Fiji^63^.

Data quantification was performed by counting cells in 3 individual fields of views for each cell line. Data are represented by mean ± SEM. Pearson correlation coefficients were determined using the colocalize plugin in Fiji^63^ by calculating for 10 individual cells per cell line.

### LDH cytotoxicity assay

WT iBMDMs, NLRP3^-/-^ iBMDMs and NLRP3^-/-^ iBMDMs reconstituted with human WT and mutant mScarlet-NLRP3 were seeded at 2.5×10^6^ cells/well on a 96-well tissue culture plate overnight. On the next day the cells were treated with 1 μg/ml LPS (Invivogen, Cat. no: tlrl-b5lps) for 3.5 h or with 1 μg/ml LPS for 2.5 h and additionally with 20 μM MCC950 (Millipore Sigma, Cat. no: 538120) for 1h followed or not by NLRP3 activation with 20 μM nigericin (Sigma-Aldrich, Cat. no: N7143-5MG) for 1.5 h. Cell supernatants were analyzed for LDH activity using the CytoTox 96® Non-Radioactive Cytotoxicity Assay Kit (Promega, Cat. no: G1780).

### Data availability

The cryo-EM maps have been deposited in the Electron Microscopy Data Resource under the accession numbers EMD-23302 (6-fold NLRP3 double ring cage purified in the presence of dATP) and EMD-23303, EMD-23304, and EMD-23305 (6-, 7- and 8-fold NLRP3 double ring cages, respectively, purified in the presence of ADP and MCC950). The atomic coordinates have been deposited in the Protein Data Bank under the accession number 7LFH (6-fold NLRP3 cage purified in the presence of dATP). All constructs and additional information can be obtained from the corresponding author upon reasonable request.

